# *Nostoc* talks back: Temporal patterns of differential gene expression during establishment of the *Anthoceros-Nostoc* symbiosis

**DOI:** 10.1101/2021.10.27.465970

**Authors:** Poulami Chatterjee, Peter Schafran, Fay-Wei Li, John C Meeks

**Affiliations:** Department of Microbiology and Molecular Genetics, University of California, Davis, CA 95616; Boyce Thompson Institute, Ithaca, NY, USA; Plant Biology Section, Cornell University, Ithaca, NY, USA

**Keywords:** Hornwort *Anthoceros punctatus*, Cyanobacterium *Nostoc punctiforme*, Nitrogen fixation, Nitrogen starvation, RNA sequencing, Symbiosis, Time course

## Abstract

Endosymbiotic association between hornworts and dinitrogen-fixing cyanobacteria form when the plant is limited for combined nitrogen (N). We generated RNA-Seq data to examine the temporal gene expression patterns during culture of N-starved *Anthoceros punctatus* in the absence and presence of the symbiotically competent cyanobacterium *Nostoc punctiforme*. Symbiotic nitrogenase activity commenced within 5 days of coculture reaching a maximal by 14 days. In symbiont-free gametophytes, chlorophyll content, chlorophyll fluorescence characteristics and transcription of genes encoding light harvesting and reaction center proteins, as well as the small subunit of ribulose-bisphosphate-carboxylase/oxygenase, were downregulated. The downregulation was complemented in a temporal pattern corresponding to the *N*. *punctiforme* provision of N_2_-derived ammonium. The impairment and complementation of photosynthesis was the most distinctive response of *A*. *punctatus* to N-starvation. Increases in transcription of ammonium and nitrate transporters and their *N*. *punctiforme*-dependent complementation was also observed. The temporal patterns of differential gene expression indicated *N*. *punctiforme* transmits signals to *A*. *punctatus* both prior to, and after its provision of fixed N. This is the only known temporal transcriptomic study during establishment of a symbiotic nitrogen-fixing association in this monophyletic evolutionary lineage of land plants.

**Highlights:** Temporal RNA-Seq analysis revealed how symbiotic cyanobacteria impact plant partners’ global gene expression and elucidated the nature of bidirectional communications between the partners

## Introduction

Nitrogen (N) is an essential nutrient for all life on earth and is the most limiting nutritional factor governing crop productivity around the world (Guo *et al.*, 2019). N-limitation results in metabolic instability, which in oxygenic photosynthetic organisms is manifest by light-dependent reductant accumulation leading to the generation of toxic reactive oxygen species.

Nitrogen (N) is an essential nutrient for all life on earth and is the most limiting nutritional factor governing crop productivity around the world (Guo *et al*., 2019). N-limitation results in metabolic instability, which in oxygenic photosynthetic organisms is manifest by light-dependent reductant accumulation leading to the generation of toxic reactive oxygen species.

As plants are sedentary, they are dependent on the available nutrients present in their rhizosphere. They acquire, metabolize, and recycle to maintain their nutrient availability throughout their life course (Pérez-Jaramillo *et al*., 2016). Plants perceive and respond to the stress of N deficiency via numerous physiological and metabolic events (Hsieh *et al*., 2018), including enhanced transcription of nitrate and ammonium transporters (Crawford and Glass, 1998; Glass, 2002) and ubiquitination of proteins, leading to their turnover and N recycling (Liu *et al.,* 2017). The CO_2_ fixing enzyme, ribulose-bisphosphate-carboxylase/oxygenase (RuBisCO) contributes around 50% of the protein content in the leaves and is one of the main reservoirs of nitrogen (Feller *et al*., 2008). The N-starvation dependent downregulation of transcription of plant genes leads to the depletion of RuBisCO, chlorophyll-binding proteins of the light harvesting complexes, and photosynthetic electron transport (Logan *et al*., 1999; Paul and Driscoll, 1997).

A few lineages of plants have acquired the ability to establish symbiotic associations with nitrogen-fixing bacteria, allowing them to colonize N-poor habitats. Because the nitrogenase enzyme complex is highly sensitive to oxygen inactivation, plants in these associations must have evolved accommodations for oxygen protection. The most well-known and studied is associations between leguminous plants and rhizobia bacteria, the latter of which have an obligate respiratory energy metabolism. Synthesis of legume hemoglobin (leghemoglobin), through cooperation of both symbiotic partners, modulates oxygen tension in the root nodules (O’Brian, 1996). Filamentous aerobic actinobacteria, such as *Frankia* spp., fix nitrogen in free-living and symbiotic growth states in the oxygen-protective, hopanoid lipid-enriched vesicles at the tips of primary and secondary filaments (Ghodhbane-Gtari *et al*., 2014). Certain filamentous cyanobacteria fix nitrogen in specialized cells called heterocysts, which produce a bilayered glycolipid and polysaccharide outer wall that effectively restricts solute and gas diffusion (Walsby, 2007). Heterocyst-forming cyanobacteria, mainly of the genus *Nostoc*, establish symbiotic associations with specific representatives of four of the five major divisions of land plants. Only in association with the angiosperm *Gunnera* spp. is the *Nostoc* symbiont intracellular in specialized glands at the base of the petiole. In associations with gametophytes of hornworts and two liverworts, leaves of the water fern *Azolla* spp. and coralloid (secondary) roots of cycads, the *Nostoc* spp. are endophytic, but extracellular in specialized preformed cavities or layers (Adams, 2002).

Land plants evolved from a charophycean algal ancestor approximately 470-515 million years ago, and consist of two monophyletic lineages: tracheophytes (vascular plants) and bryophytes (i.e., mosses, liverworts, and hornworts) (Szövényi *et al*., 2021). Bryophytes are, perhaps, the oldest lineage of land plants to form stable long-lasting endosymbiotic associations with nitrogen-fixing bacteria, in this case oxygenic photoautotrophic cyanobacteria (Sprent and Raven, 1985). We have utilized pure cultures of the hornwort *Anthoceros punctatus* and the model symbiotic cyanobacterium *Nostoc punctiforme* strain American Type Culture Collection 29133 (syn Pasteur Culture Collection 73102) (hereafter *N. punctiforme*) as an experimental system. Some form of motility in at least one partner is required for efficient establishment of a symbiosis. *Nostoc* spp. vegetative filaments are nonmotile and motility requires the transient differentiation of motile-by-gliding filaments called hormogonia; hormogonium filaments lack heterocysts, do not fix nitrogen and are growth-arrested (Tandeau de Marsac, 1994). N-limited *A. punctatus* produces an extracellular compound termed a hormogonium inducing factor (HIF) that induces synchronous differentiation of hormogonia (Campbell and Meeks, 1989) and, most likely, chemoattractants (Nilsson *et al.,* 2006). Once hormogonia have colonized slime cavities on the ventral surface of the gametophyte, further hormogonium differentiation is suppressed by a plant produced hormogonium repressing factor and the hormogonia return to the vegetative growth state (Cohen and Meeks, 1997). Under these conditions, infection is confined to the initial 2-3 days of coculture. Colonization of the slime cavity is followed by growth of associated *N. punctiforme*, concurrent heterocyst differentiation to a level that is about 3-fold higher than in the free-living state and a rate of nitrogen fixation that is more than 8-fold greater (Meeks, 1998, 2003). The fixed nitrogen is released as ammonium to support growth of *A. punctatus* (Meeks *et al*., 1985).

Molecular mechanisms of infection and organogenesis have been identified by genetic analyses and modeled in rhizobia-legume and actinobacteria-actinorhizal plants in the context of a common symbiotic signaling pathway (CSSP), collectively with arbuscular mycorrhizal fungi and nearly all land plants (Horváth *et al*., 2011; Harris *et al*., 2020). Apart from a snapshot RNA-Seq experiment during the genome sequencing of the hornworts *A. punctatus* and two strains of *A*. *agrestis* (Li *et al*., 2020), nothing is known of the genetics or genomics of N-starvation dependent infection by symbiotically competent *Nostoc* species of any plant association.

Since N-starvation is a prerequisite for endophytic symbiotic association in plants, we initiated a time course RNA-Seq analysis of *A*. *punctatus* gametophyte tissue incubated in the absence of combined N and absence or presence of *N*. *punctiforme*. We anticipate that the results will resolve the identity of differentially expressed genes (DEG) and possibly the metabolic pathways altered in symbiotically associated *A*. *punctatus* and provide experimental approaches to identify the controlling regulatory elements. How a hornwort responds to N-starvation and initiates symbiotic association can then be compared to other nitrogen-fixing symbioses and inform approaches to engineering similar symbioses in crop plants (Mus *et al*., 2016; Oldroyd, 2013; Pankievicz *et al*., 2019).

Indeed, the results show that the presence of *N*. *punctiforme* does alter the patterns of DEG, relative to N-starvation of *A*. *punctatus* alone, by enhancing transcription of some genes that were downregulated or by repressing genes that were upregulated. Analysis of the temporal patterns of transcript accumulation indicates that complementation of about 42% of the DEG by *N*. *punctiforme* correlated with the onset of symbiotic N_2_ fixation, while about 25% of the DEG were complemented during colonization of the symbiotic cavity before N_2_ fixation had commenced.

## Materials and Methods

### Plant and Cyanobacterium Growth Conditions

Previously, surface-sterilized spores were germinated to obtain axenic gametophyte tissues of *A. punctatus* (Enderlin and Meeks, 1983). Hutner’s minimal medium with NH_4_NO_3_ as the nitrogen source (H+N) was used for the growth of *A. punctatus* (Enderlin and Meeks, 1983). The minimal medium was supplemented with 5 mM 2-(N-morpholino) ethanesulfonic acid (Mes: Sigma Chemical Co.) adjusted to pH 6.4 with NaOH as buffer and 0.5% (w/v) glucose, originally included to increase the growth rate of light-limited laboratory cultures. Gametophyte tissues were incubated in 100 ml of medium in 300 ml Erlenmeyer flasks, at 20° C and 50 rpm orbital shaking, under 20-Watt fluorescent lamps (3.5-8.0 W m^−2^; cool-white) with 16 h and 8 h of the light-dark cycle. The stock cultures of symbiont-free *A. punctatus* were subcultured into H+N plus Mes and glucose every 14 days.

*N. punctiforme* was maintained under standard growth conditions (Enderlin and Meeks, 1983). For experiments, the cultures were grown up to an early light-limited linear phase in the standard minimal salts medium diluted fourfold, with N_2_ as the nitrogen source (Enderlin & Meeks, 1983). The cultures were incubated in 50 ml of medium in a 125 ml Erlenmeyer flask with orbital shaking at 120 rpm and 25° C under 19 to 46 W m^−2^ s^−1^ of cool white fluorescence lights. Chlorophyll *a* (Chl) content of *N. punctiforme* was determined after extraction in 90% methanol (Meeks and Castenholz, 1971).

### *A. punctatus*-*N. punctiforme* Symbiotic Reconstitution and Sampling

Gametophyte tissues of *A. punctatus*, grown in H+N were washed with Hutner’s medium lacking combined N (H-N) and transferred to two 100 ml of H-N medium, with Mes and glucose (in 300 ml Erlenmeyer flasks). Approximately 10-12 g fresh weight (FW) of *A. punctatus* gametophytes (14 days after the last transfer) were added to each of the flasks, one was cocultured with 100-135 μg Chl *a* content of *N. punctiforme*, whereas the other flask was symbiont-free. The flasks were incubated under the growth conditions noted above. The gametophytes were sampled at day 0 (before initiation of the N-starvation), and after 2, 5, 7, 10, 14, and 28 days of incubation. The whole experiment was replicated thrice, with a total of 39 samples.

### Acetylene Reduction Assay, Total Chl estimation, and Chl Fluorescence Measurements

Nitrogenase activity was measured by the reduction of acetylene to ethylene. Approximately 100 mg of tissue was sampled and washed using streams of distilled water to remove epiphytic attachment of *N. punctiforme* (as were all cocultured tissues prior to experimental analysis). The cleaned sample was incubated in 2 ml of H-N medium in a 7.5 ml glass vial sealed with a septum stopper. Acetylene was made freshly from calcium carbide (CaC_2_) and injected into each vial to 6% (vol/vol). The ethylene-acetylene content was monitored by gas chromatography after 30, 60, and 90 min of incubation. One hundred microliter of a sample from the vial atmosphere was injected onto a Porapak R column in a gas chromatograph equipped with a flame ionization detector (model 940, Varian Instrument Division, Walnut Creek, CA). The normalization and calculation of the rate of ethylene production using the excess acetylene as an internal standard (i.e., ratio of ethylene to acetylene equals amount of ethylene produced based on a standard curve) were performed according to Steinberg & Meeks (1991). The number of *N. punctiforme* colonies in cocultured gametophytes was determined under a stereomicroscope.

The total Chl *a* and Chl *b* content from N-starved cocultured and symbiont-free gametophytes was estimated after extraction of 100 mg (freshly ground sample in liquid nitrogen [LN]) of plant sample in 10 ml of 80% Acetone and quantified using the dichromatic equations of Inskeep & Bloom (1985).

Dark-adapted (20 min darkening) Chl fluorescence yield, (*F*_v_/*F*_m_) was estimated with an imaging PAM fluorometer (PAM-2000, Walz, Effeltrich, Germany) with a saturating pulse (7000 μmol m^−2^ s^−1^ for 1 s) of blue light to measure the maximum dark-adapted fluorescence yield, *F*_m_. The maximum and effective quantum yields of Photosystem II electron transport were calculated as *F*_v_/*F*_m_ =◻(*F*_m_ −*F*_0_)/ *F*_m_.

All the above-mentioned experiments were repeated thrice (biological replicates), and three technical replicates for each time point.

### Improved *A. punctatus* Genome Assembly and Annotation

Following updates to the Oxford Nanopore’s Guppy basecaller, the raw sequence data used to assemble the first iteration of the *A. punctatus* genome (Li *et al.*, 2020) was re-basecalled and assembled with Flye (Kolmogorov *et al.*, 2019). The draft assembly was polished with five iterations of Pilon (Walker *et al.*, 2014) using 100X coverage. A custom repeat library was created with EDTA (Ou *et al.*, 2019) and input into RepeatMasker (Smit *et al.,* 2015) to mask repetitive regions of the genome for gene annotation. Gene models were predicted with BRAKER2 (Brůna *et al.*, 2021) using RNA-seq data from one replicate of the Li *et al.* (2020) cyanobacterial symbiosis experiment, as well as their predicted proteins from *A. agrestis* as training data. Gene functional annotations were obtained using eggNOG-mapper (Cantalapiedra *et al.*, 2021). Genome annotation completeness was measured using BUSCO v5 with the Viridiplantae odb10 reference dataset (Simão *et al.*, 2015).

### Plant RNA Extraction and RNA Sequencing Protocol

The freshly sampled and washed gametophytes were ground in LN with a mortar and pestle and stored at −80° C. RNA was extracted from 100 mg of frozen tissue using the Spectrum total RNA plant kit (Sigma-Aldrich). The mRNA library was prepared by poly-A enrichment and paired-end sequencing was performed on Illumina NovaSeq6000 (2 × 150 bp) by Novogene (Sacramento, CA).

### RNA-Seq Data Set and Sequencing Analysis of *A. punctatus*-*N. punctiforme* Symbiosis

All fastq files were quality-filtered and trimmed using fastp version 0.20.0 (Chen *et al.*, 2018*b*) and subsequently mapped to the reference genome using HISAT2 version 2.1.0 (Kim *et al.*, 2015). All reads were 96.33% to 98.57% mapped to the genome. Transcript abundances were estimated using the updated *A. punctatus* genome annotation in Stringtie version 2.1.1 (Pertea *et al.*, 2015).

The median of ratios method of normalization was used to normalize all the gene counts, which were later used to plot graphs of different temporal gene expression patterns from selected clusters and a few other gene expression patterns from the transcriptome. The functional annotation file containing all KEGG and GO annotations for 69% of the transcripts from the gene count matrix are detailed in Supplemental Dataset S1. Variance stabilizing transformation (VST) normalized genes were fed into maSigPro (Conesa *et al.*, 2006) in R version 4.0.2 to analyze the changes in the temporal gene expression patterns. The significantly (*P* ≤ 0.05) DEG were subsequently clustered into nine different profiles as per their gene expression patterns using “hclust” in maSigPro.

### Statistical Analysis

The results of cocultured or symbiont-free *A. punctatus* gametophytes in acetylene reduction assays, total Chl content, and *F*_v_/*F*_m_ were tested using ANOVA followed by Tukey HSD test with the function aov and tukeyhsd which is a part of R core package stats, version 4.0.2 (Yandell, 2017).

## Results and Discussion

### Time Course of *N. punctiforme* Colonization of *A. punctatus*, Induction of Nitrogenase Activity and Change in Chlorophyll Content in N-Starved Gametophyte Tissue

The colonization of *N. punctiforme* within the slime cavities of *A. punctatus* gametophyte tissue is the third step of the infection process, following plant-dependent induction of hormogonium differentiation and chemoattraction of hormogonia. *N. punctiforme* colonies were routinely microscopically visible in gametophytes after 7 days of co-culture and only rarely observed earlier (Fig. 1A). The colonies were largely present at the tip of the gametophytes, near the marginal meristem. The color density of the day 7 colonies was relatively light, but it increased with time up to 14 days after coculture, reflecting growth of symbiotic *N. punctiforme* in the slime cavity. The space between the gametophyte margin and the colonies increased as the tissue also continued to grow (Fig. 1A), but no new infections, normalized to fresh weight (FW) of gametophyte tissue, were observed over the 28-day period. The reconstituted *A. punctatus-N. punctiforme* tissue retained its green color throughout the experimental time course. Conversely, when symbiont-free *A. punctatus* was N-starved, the gametophytes appeared pale green on day 2, then turned yellow to pale yellow as incubation time continued (Fig. 1B).

**Figure 1:**
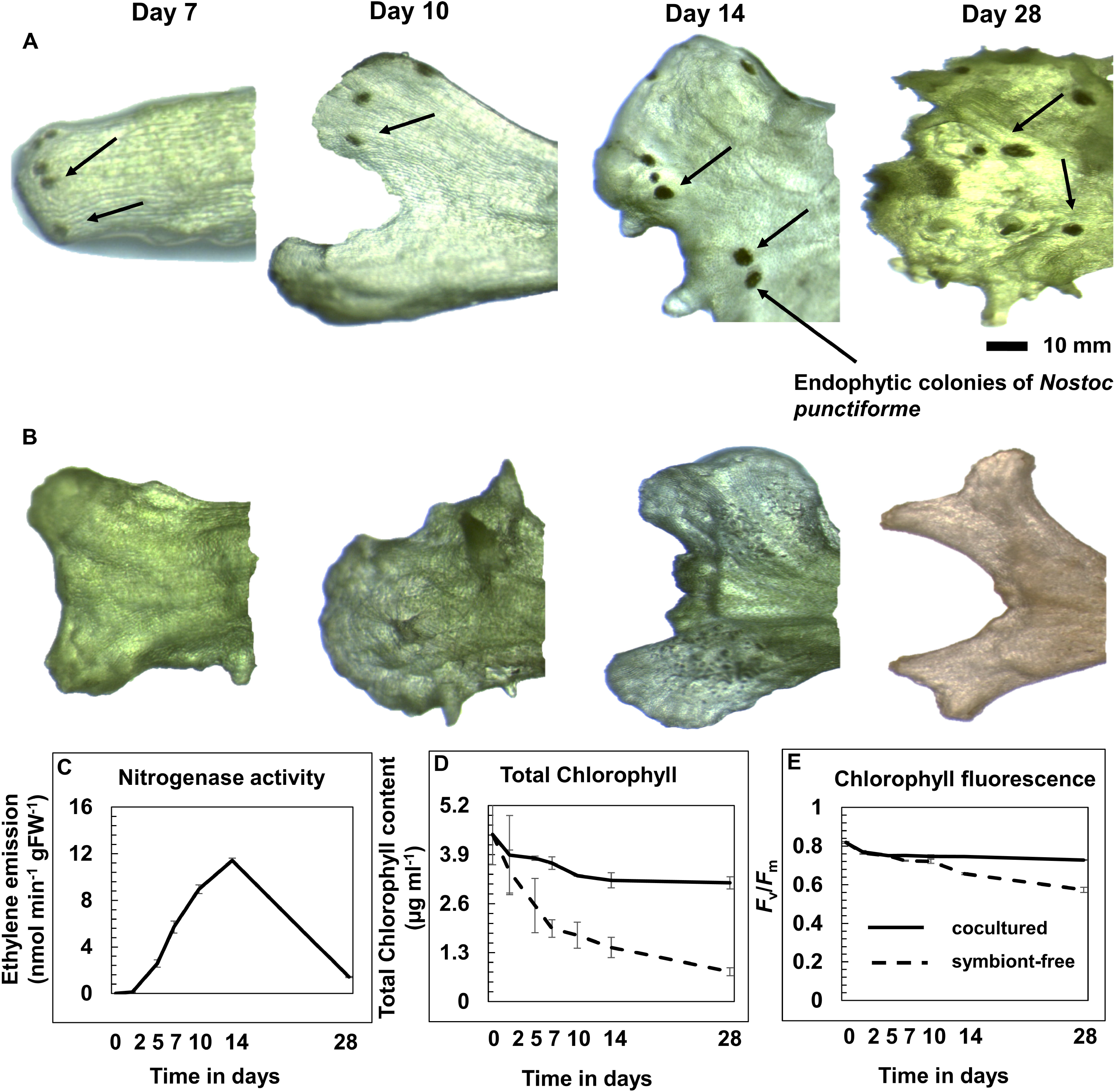
Morphological and physiological changes in symbiotic and symbiont-free gametophytes during N-starvation. Micrographs of colonization of *N. punctiforme* inside the slime cavities of *A. punctatus* gametophytes (A) and symbiont-free *A. punctatus* gametophytes (B), and their visual changes with time during N-starvation. Average rates (±SE) of acetylene reduction (nitrogenase activity) from symbiotic gametophytes (C). Average content (±SE) of total chlorophyll (D) and dark-adapted (20-minute darkening) maximum quantum yield of PS II estimated by chlorophyll fluorescence (*F*_v_/*F*_m_) (E) in the gametophytes of symbiotic and symbiont-free *A. punctatus* under N-starvation conditions.

Nitrogenase activity was monitored by the acetylene reduction assay. Acetylene conversion to ethylene was observed within 5 days of coculture, before *N. punctiforme* colonies could routinely be microscopically observed in the tissue. The rate of acetylene reduction increased about 4-fold between at days 5 and 14, and then decreased to 12% of the maximal rate by day 28 (P < 0.001) (Fig. 1C). *N. punctiforme* does not form a stable and long-lasting association with *A. punctatus* and this is reflected, in part, by the decline in acetylene reduction 14 d after the initiation of coculture. We use *N. punctatus* as a model organism to examine the initial stages of symbiotic association because it is amenable to facile genetic manipulation (Cohen *et al*., 1994; 1998), in contrast to an original stable symbiotic isolate from *A. punctatus*, *Nostoc* sp. strain PCC 9305 (syn strain UCD 7801). The rate of ethylene production from symbiotic gametophytes increased with time from 0.109 to 11.423 nmol min^−1^ g FW^−1^ (Fig. 1C). The day 14 rate is similar to that of *A*. *punctatus* colonized with *Nostoc* sp. strain PCC 9305 (Steinberg and Meeks, 1991).

Total Chl content of N-starved cocultured and symbiont-free *A. punctatus* was estimated by calculating the Chl *a* and Chl *b* content from gametophyte tissues. The total Chl content in symbiotic gametophytes varied between 5.23 to 3.3 μg per 10 mg FW, whereas the total Chl content declined from 5.23 to 0.79 μg per 10 mg FW in symbiont-free *A*. *punctatus* (P < 0.001) (Fig. 1D).

### Improved Genome Assembly and Annotation of *A. punctatus*

The new genome assembly increased in size slightly from 132.8 to 134.6 Mb and increased in contiguity substantially, as contig N50 doubled from 1.7 to 3.3 Mb. The number of predicted gene models from the new assembly decreased from 25,426 to 23,021 but there was a considerable increase from 85% to 92% complete genes identified by BUSCO (Dataset S1). Of all the predicted genes, 69% can be functionally annotated (Dataset S2). The gene location numbers are formally assigned as, for example: Anthoceros_punctatus_v2_contig1_g00020; on occasion, we refer to the location numbers in figures or text as, for examples, contig1_g00020 or g00020.

### Global RNA-Seq Analysis

A two-component Principal Component Analysis (PCA) identified the variation in the biological replicates at each time point for each treatment. Overall, all the biological replicates are closely associated with each other. The exception is the three biological replicates at time 0, which are less tightly clustered (Fig. S1).

### Identification of Differentially Expressed Genes by Symbiotic and Symbiont-Free *A. punctatus*

Analysis by maSigPro yielded 1,448 DEG of *A*. *punctatus* at a P < 0.05 through the two six-point time courses without and with *N. punctiforme*, although the biochemical functions of only 1,210 transcripts (84%) with annotations were predicted via GO and/or KEGG (Dataset S3). This 84% is markedly higher than the 69% of genes that were annotated in the genome (Dataset S2). The plots of both experimental lines include counts from the common time 0.

A user selected output of 9 clusters yielded multiple cluster patterns reflecting initial up- or downregulation of gene expression during the time course of N-starvation, persistence of the relative amount of transcript accumulation with time and a time dependence upon reaching the stable or final relative value of transcript accumulation. In the case of symbiont-free *A*. *punctatus*, six temporal patterns can be described (Fig. 2). **i)** Two clusters (1 and 6) in which genes whose transcript accumulation was upregulated to different levels and at different rates between times 0 and 5 d, and the level of transcripts remained at an elevated value, relative to time 0, for up to 28 days. **ii)** Two clusters (4 and 9) where transcripts were rapidly upregulated, followed, or not, by a slight delay before declining to approximately the time 0 value by day 28. **iii)** In cluster 8, which also represents the fewest genes, transcript accumulation was upregulated by day 5, remain elevated until day 10 before declining to the time 0 value by day 28. **iv)** Two clusters (3 and 7) in which transcripts rapidly declined to a lower constant level by days 2 to 5 and then remained depressed through day 28. **v)** In cluster 5, transcripts rapidly declined by day 2 and then slowly increased, approaching the time 0 level. **vi)** Transcripts in cluster 2 showed a slight delay in changes from the time 0 level before their accumulation slowly declined to a lower level.

**Figure 2:**
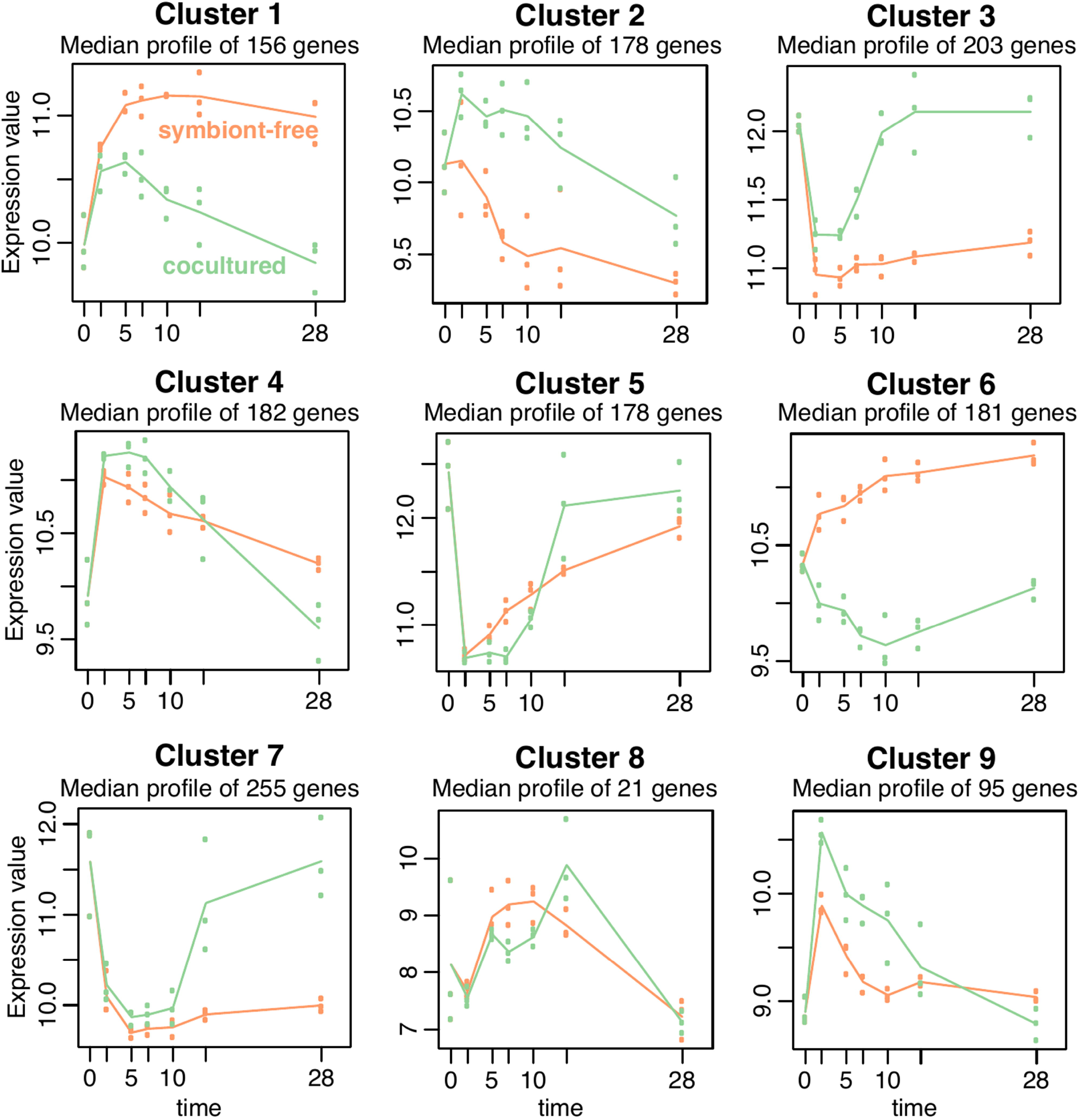
Temporal expression profiles for differentially transcribed genes from cocultured and symbiont-free *A. punctatus* gametophytes. The number of clusters was user selected and then arbitrarily numbered by the algorithm. The ordinate is the median log of the VST normalized gene expression for that set of genes (*P* ≤ 0.05). The abscissa is time in days since initiation of N-starvation. Orange and green lines indicate differential gene expression profiles of symbiont-free and cocultured gametophytes, respectively.

When *A*. *punctatus* was cocultured with *N*. *punctiforme*, allowing for reconstitution of the symbiosis, there were two overarching groups displaying patterns of transcript accumulation. In the first group, clusters 4, 8 and 9 showed essentially the same patterns in the presence or absence of *N*. *punctiforme*. The shape of the plot of the cocultured *A*. *punctatus* in cluster 5 appears similar to symbiotic tissue in clusters 3 and 7; the similarity comes from the sharp downregulation followed by a 5-day lag and then upregulation starting at day 7 before peaking at 14 days of incubation. This appears to reflect a hybrid response because there was slow upregulation in the absence of *N*. *punctiforme* and the level of expression in both tissues approaches the time 0 value by day 28. Here, we will treat the transcriptional patterns in clusters 4, 5, 8 and 9 as influenced by, but not dependent on, *N*. *punctiforme*. Most interesting is the second group; relative to *A. punctatus* alone, the presence of *N. punctiforme* unequivocally repressed the N-starvation induced upregulation of transcript accumulation (clusters 1 and 6) and enhanced the transcription of genes that were downregulated in its absence (clusters 2, 3 and 7). However, there are temporal differences in the responses. For examples, in cluster 6, the repressive effect of *N. punctiforme* appears to be immediate, whereas in cluster 1 there was an immediate transcriptional upregulation followed by a short period of stable expression and then a first order decline to the time 0 level. In cluster 2, the cocultured tissue showed an immediate upregulation in transcripts, which remained elevated until at least day 10, before beginning a decline to slightly less that the time 0 value, while that in symbiont-free *A. punctatus* stayed constant through day 2 before slowly declining to a lower level. The transcriptional patterns in clusters 3 and 7 differ in the timing of the enhanced accumulation; the transcripts in cluster 3 began to increase accumulation from the repressed level after 5 days of incubation, while in cluster 7 the increase started after day 10. In both cases, transcripts returned to at or near the time 0 level, which may indicate return to a steady state.

### Functional Analysis of Differentially Expressed Genes

The differentially expressed, annotated, transcripts were manually organized into the 10 major metabolic groupings shown in Fig. S2 to allow analysis of their expression patterns in the context of cellular growth and its regulation. The most highly represented genes are present in the unassigned function and core metabolism categories, which might be expected in global transcriptomic analysis of a growth stress response and where 56% of the DEG were immediately downregulated. With respect to its broad descriptive nature, the functional analysis is presented in Dataset S3 with a complete gene list, including normalized expression values and a guide to their interpretative navigation (Supplementary Guide S1). In the following section, we will focus on those DEG complemented by the presence of *N*. *punctiforme*, as well as some genes that were sorted as constitutively expressed but that can be predicted as relevant to N-starvation and symbiotic interactions.

### Genes Relevant to Nitrogen Starvation and Symbiotic Interaction

In the response of *A*. *punctatus* to N-starvation, based on multiple studies of algae and terrestrial plants, we hypothesize that managing the resulting excess photosynthetic reductant pool could occur by three different processes: **i)** decrease in the rate of photosynthetic reductant generation; **ii)** increases in the uptake of low levels of an existing source or acquisition of an alternative environmental source of combined N; or **iii)** utilization of an organic reductant sink whose metabolic product can be stored or excreted; or any combination of these. If any one of these three processes were to exclusively occur during N-starvation of *A*. *punctatus*, the result is predicted to be an initial up or down transcriptional-regulation of the genes encoding the essential proteins and would yield predictable temporal patterns and levels of transcript accumulation. Here, we have reconstituted the *A*. *punctatus*-*N*. *punctiforme* symbiotic association such that N_2_-derived ammonium would become the alternate environmental source of N. The time in which *N*. *punctiforme* becomes a functional N_2_-fixing symbiont after initiation of coculture was determined by whole tissue assays of acetylene reduction as a proxy of nitrogenase activity. The data in Fig. 1c indicates that *N*. *punctiforme* linearly attains a fully functional symbiotic state between shortly after the initiation of coculture at day 5 up to day 14; the nitrogenase specific activity is the same on day 14 as that of a long-term association between *A*. *punctatus* and its original symbiotic isolate, *Nostoc* sp. strain PCC 9305 (Steinberg and Meeks, 1991). Thus, although *N*. *punctiforme* does not form a long-term symbiosis with *A*. *punctatus*, its N_2_-fixing physiology, by 14 days of association, is reflective of a long-term association.

In this context, we suggest that clusters 4, 5, 8 and 9, where the transcription patterns are not markedly influenced by the presence of *N*. *punctiforme,* reflect a general stress response that is separate from, but possibly instigated by, N-starvation response; these clusters represent 33% of the DEG. Our discussion below is largely, but not exclusively, guided by genes in clusters 1, 2, 3, 6 and 7 in which the presence of *N*. *punctiforme* does alter the patterns of expression. Specifically, the temporal patterns in clusters 1, 3 and 7 strongly correlate with the analogous pattern of the onset of symbiotic nitrogenase activity and account for 42% of the DEG. Conversely, the temporal patterns in clusters 2 and 6 are initiated prior to, and appear independent of, symbiotic dinitrogen fixation; these clusters account for 25% of the DEG. We will also introduce some constitutively expressed genes encoding proteins that could be predicted as involved in adaptation to these responses. In tracheophytes, gene products and isozymes are often localized in specific tissues and organs, which help to understand the physiological roles of gene family members. Those bryophytes that establish symbioses may also have tissue differentiated from the bulk gametophyte thallus, such as sporophytes, rhizoids and slime cavity cells, which, in the liverwort *Blasia pusilla*, elaborate septate, branched filaments of metabolite transfer cells that increase in profusion after *Nostoc* spp. colonization (Rodgers and Stewart, 1977), and where isozymes could be localized.

### Genes Related to the Consequences of N-Starvation on Photoautotrophic Growth of *A*. *punctatus* and Their Transcriptional Complementation by *N*. *punctiforme*

#### Photosynthetic characteristics

Fig. 1B depicts the visual degreening of symbiont-free gametophyte tissues in absence of combined N, which corresponds with the lowering of total Chl content in those tissues, as seen in other plants (Sayed, 1998). Nitrogen limitation in chloroplasts impairs the photosynthetic machinery, causing significant damage in the reaction centers of photosystem (PS) II, leading to a major reduction in *F*_v_/*F*_m_ (Zhao *et al.,* 2017) (Fig. 1E). The maximum dark-adapted PS II quantum yield was estimated by in vivo Chl fluorescence (*F*_v_/*F*_m_). A sustained *F*_v_/*F*_m_ value, ranging from 0.729 to 0.819 was observed in symbiotic *A. punctatus* gametophytes. N-starvation reduced *F*_v_/*F*_m_ of symbiont-free *A. punctatus* from 0.819 at day 0 to 0.574 (stressed phase) by day 28 (P < 0.001). It should be noted that a much lower stressed phase value than the 0.517 at day 28 has been reported in plants under different stress conditions (Murchie and Lawson, 2013). The latter comparison indicates photosynthesis was not completely switched off in symbiont-free *A*. *punctatus*, although visually the tissue looked compromised; this conclusion is supported by the continued transcription of genes in, for example clusters 1 and 6 by day 28.

There are 33 DEG encoding proteins indirectly or directly related to photosynthesis (Dataset S3). Sixty percent of the chloroplast and photosynthetic transcripts are present in cluster 3 in which they were initially downregulated and then upregulated in the presence of *N. punctiforme* in a temporal pattern corresponding the production of N_2_-derived ammonium. Examples of transcripts encoding one of ten related or homologous proteins in a LHC (LHCA2-2, associated with PS I); a PS II reaction center protein (PSBW, which stabilizes macromolecular complexes) and one of two oxygen-evolving enhancer proteins (PSBP, which are also required for PS II core stability); a PS I reaction center protein (PSAD, probable ferredoxin docking protein); and the RuBisCO small subunit are shown in Fig. 3A-E. The transcription of these core photosynthetic genes was rapidly downregulated and appears to be uncoupled from the slower decline in total Chl content and even slower decline in PS II quantum yield. Some of these internal differences might be due to the relatively low light intensity (ca. 65 μmol m^−2^ s^−1^) used during culture, which could slow bleaching and photooxidation. This suggestion is supported, in part, by the observation that a gene encoding a protein involved in xanthophyll metabolism is present in cluster 7 where it showed rapid downregulation in symbiont-free and symbiotic tissues and was subsequently upregulated in symbiotic tissue between days 14 and 28 to the approximately the time 0 level (Fig. 3F). This pattern implies a degree of N-starvation response, although it is the reverse of that anticipated in protection from a highly reduced photosynthetic electron transport system. Xanthophylls are present in association with Chls in the LHCs where they operate in a cycle of oxidation and reduction to quench excitation generated by high light intensities (Niyogi *et al.*, 1997; Latowski *et al.*, 2011). Genes encoding core components of the PS reaction centers and of electron transport chains, as well as the large subunit of RuBisCo, could also be differentially transcribed, but they are localized in the plastid genome and their transcription proposed to be subject to redox control (Allen, 2015); thus, we did not sequence them due to the lack of poly-A extensions.

**Figure 3:**
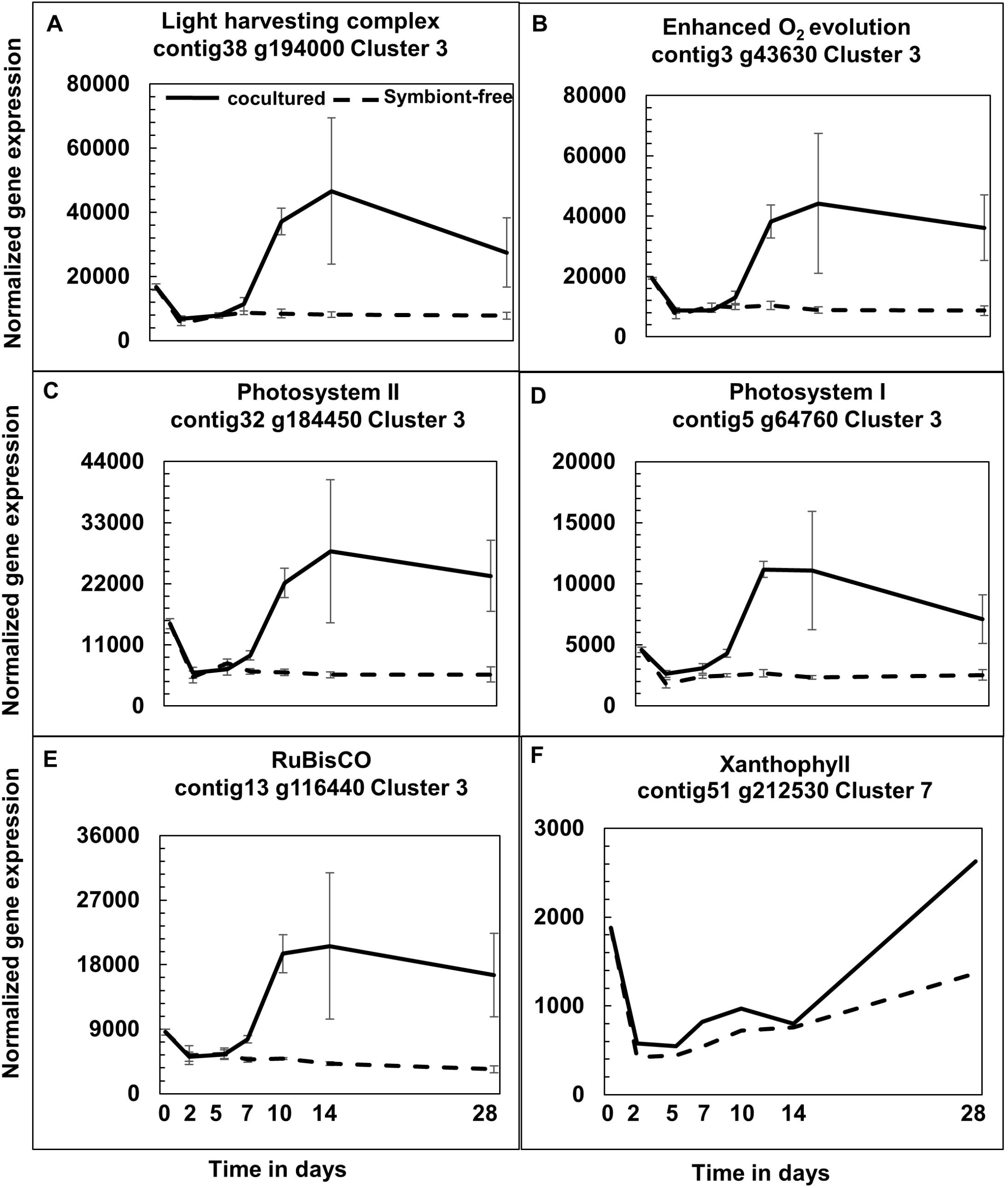
Temporal patterns of normalized expression of genes encoding proteins involved in photosynthesis. The ordinate is the mean (±SE) normalized gene expression of transcripts (*P* < 0.05). The differentially expressed genes encode proteins for: a representative light harvesting complex (A), enhanced O_2_ evolution (B), PS II reaction centers (C), PS I reaction centers (D), RuBisCO small subunit (E), and xanthophyll metabolism (F), from cocultured and symbiont-free *A. punctatus* gametophytes. Time is in days as in Fig. 2.

We conclude that a major physiological and transcriptional response to N-starvation by *A*. *punctatus* was to lower the photosynthetic potential, largely by destabilizing the PS II reaction center complex and limiting electron transfer out of the PS 1 complex. The lowered potential was subsequently complemented by symbiotic association with N_2_-fixing *N*. *punctiforme*.

#### N acquisition and assimilation

The genes encoding N acquisition and assimilation in *A*. *punctatus* occur in multigene families; they are summarized in Supplementary Table S1. Some N-starved plants induce the transcription of transport systems for ammonium or nitrate/nitrite (Krapp *et al.*, 2011; Calabrese *et al.*, 2017). The *A*. *punctatus* genome contains 7 genes annotated as ammonium transporters (AMT). Redundancy in AMT genes is common in land plants, (Loqué and von Wirén, 2004; Couturier *et al.*, 2007). The genes have distinct phylogenies that can be subdivided into subfamilies and clades, and the proteins are distributed amongst various tissues and organelles. The bryophyte liverwort *Marchantia polymorpha*, has 5 AMTs in the AMT1 clade (McDonald and Ward, 2016). Only two of the *A*. *punctatus* AMT genes can be assigned biologically significant roles in ammonium transport: differentially transcribed g54810 encoding an AMT1;5 and g162610 encoding a constitutively expressed AMT2;1 (Table S1). The transcriptional pattern of g54810 is of cluster 1 showing upregulation in *A*. *punctatus* early in N-starvation, the transcription level remained high in the absence, but slowly declined in the presence, of *N*. *punctiforme* (Fig. 4A). This temporal pattern is consistent with a search for exogenous ammonium, realized when N_2_-derived ammonium became available.

**Figure 4:**
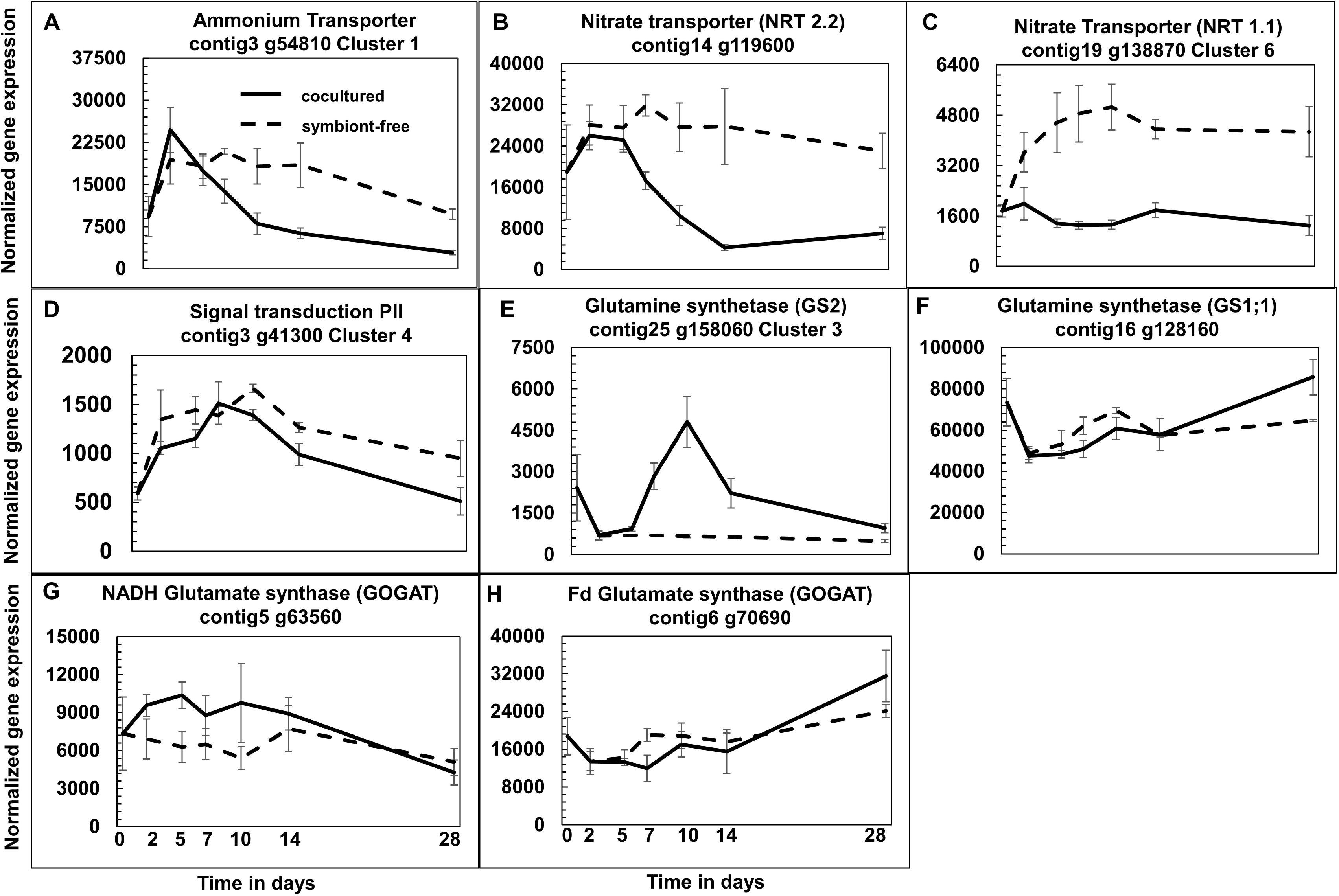
Temporal patterns of normalized expression for genes involved in N acquisition and assimilation. The ordinate is the mean (±SE) normalized gene expression of transcripts (*P* < 0.05). The selected genes encode: an ammonium transporter (A), a nitrate transporter (NRT 2.2), (B) a nitrate transporter (NRT 1.1) (C), PII signal transduction (D), a plasmid localized glutamine synthetase (GS2) (E), a cytosolic glutamine synthetase (GS1;1) (F), the expressed NADH glutamate synthase (G), and the complete Fd glutamate synthase (H), from cocultured and symbiont-free gametophytes. Time is in days as in Fig. 2.

There are 2 genes encoding nitrate transporters in the NTR2.2 family (g128250 and g119600); both were highly expressed and sorted as constitutively transcribed (Table S1). However, on inspection, both were differentially expressed in the pattern of cluster 1 (Fig. 4B). We have no facile explanation of why the algorithm did not sort these as DEGs. Nevertheless, g128250 encodes a protein with a putative signal peptide, but no transmembrane domains in the designated transport domain, thus, its actual function is not clear. The g11960 transcriptional pattern is similar AMT1:5 and consistent with a search for combined N before complementation by nitrogenase activity. The genome also contains 34 genes encoding proteins of the low or variable affinity nitrate transporters in the NTR1.1 PTR family (Sun and Zheng, 2015), now referred to as NPF (Wang *et al.*, 2018). Due to the uncertainty of their contribution to nitrate transport, this family was not thoroughly analyzed. All genes were transcribed to varying degrees ranging from 2 to 25,000 normalized counts. Four of the genes are in the differential transcriptome and present in clusters 2 (2 genes), 6 and 7. The gene in cluster 6 shows early upregulation in the absence of *N*. *punctiforme* and its upregulation was repressed in symbiotic tissue before provision of N_2_-derived ammonium (Fig. 4C).

It has been documented plants can use a variety of exogenous amino acids and even proteins as N sources for growth (Miller *et al.*, 2008; Paungfoo-Lonhienne *et al.*, 2008; Mertz *et al.*, 2019). Amino acid and other organic N transporters were present in the differential transcriptome; the transporters with the highest levels of expression are for lysine and histidine and they are present in cluster 7 (Dataset S3), which is inconsistent with N-limited complementation function. It remains possible that the presence of exogenous amino acids that, when coupled with N-starvation, could induce the transcription of their respective transporters, but we have no evidence for such a possibility. Taken together, we conclude that a search for alternative inorganic N sources, such as ammonium and nitrate, is supported by the data presented.

In general, N-starvation also induces upregulation of N signaling and assimilatory proteins. Symbiotic *A*. *punctatus* assimilates exogenous and N_2_-derived ammonium by the glutamine synthetase (GS) and glutamate synthase (GOGAT, an acronym for glutamine: 2-oxo-glutarate amido transferase) pathway (Meeks *et al.*, 1983, 1985). The GLB1 nuclear gene encodes the N regulatory protein PII, which is localized in chloroplasts. PII can be reversibly modified by uridylylation and deuridylylation and the different forms modulate both transcription and catalytic activity of target proteins, such as GS. In *A*. *thaliana*, GLB1 is transcriptionally upregulated by light and sucrose; it is downregulated in the dark and the presence of asparagine, glutamine, and glutamate (Hsieh et al., 1998). Conversely, in the green alga *Chlamydomonas reinhardtii*, GLB1 is upregulated in the absence of ammonium (Zalutskaya *et al.*, 2018). A single *A*. *punctatus* GLB1 gene encodes PII. GLB1 is rapidly upregulated during N-starvation with stable accumulation prior to declining after day 10 in the absence and presence of *N*. *punctiforme* (Fig. 4D); this pattern could reflect a modest amount of upregulation. However, the fact that the temporal DEG pattern of GLB1 transcription was not complemented by N_2_-derived ammonium implies that PII may be involved in more than N stress (Chellamuthu *et al.*, 2013).

In bacteria and some plants, limitation for ammonium results in upregulation of GS activity and synthesis, both as signaled by the covalently modified PII protein. In algae and plants, there are two forms of GS proteins; GS1, is localized in the cytosol and isozymes of it are distributed in various tissues and organs, whereas GS2 is targeted to the plastids. The plastid protein is typically encoded by a single nuclear gene, while 3 to 5 genes encode the GS1 isozymes (Swarbreck *et al.*, 2011). Eight genes are annotated as members of the GS (*glnA*) superfamily in *A*. *punctatus* (Table S1). The two primary GS genes are g158060 encoding a GS2 type of protein with a signal peptide (Fig. 4E), while constitutive expressed g162610 encodes a GS1 type protein with no signal peptide and a defined cytosolic domain (Fig. 4F). It is of interest that three other constitutively transcribed genes encode named GS proteins with signal peptides g141120 and g45770 plus g103610 which is a fusion protein with a different catalytic domain (Table S1).

Differentially expressed g158060, encoding GS2, is represented in cluster 3 where its transcripts initially declined in symbiont-free and symbiotic *A. punctatus* before they increased to a maximum by 10 d of incubation in symbiotic tissue and then again declined (Fig 4E). The pattern implies decreased transcription was in response to the absence of combined N and the subsequent increased transcription was in response to the presence of N_2_-derived ammonium. Based on the expression levels, GS1;1 would appear to be the dominant GS assimilatory protein in the entirety of the gametophyte tissue. It is not clear whether the GS2 protein is localized in chloroplasts in the bulk of gametophyte tissue, potentially at a lower concentration than GS1;1, or primarily in the slime cavity colonized by *N*. *punctiforme* as these are the first cells to encounter N_2_-derived ammonium. The post-translational modification status of any of the GS proteins in *A*. *punctatus* is unknown.

GOGAT proteins are localized to the plastids in plants (Suzuki and Knaff, 2005). There are two genes each annotated as encoding ferredoxin (Fd) (GLSF or GLU1) and NADH (GLTB) dependent GOGAT proteins in the *A. punctatus* genome (Table S1). The Fd-GOGAT encoding genes (g70690 and g70700) are present in tandem in the genome and are constitutively expressed. However, g70700 encodes a 329 amino acid peptide with only a GATase motif and a signal peptide; whether this peptide is a subunit of a holoenzyme with the g70690 protein is not clear. The g70690 gene is the most highly expressed (Fig. 4H) and most likely encodes the primary GOGAT activity in the gametophyte thallus. The gene g63560 encodes a NADH-GOGAT that consists of the core GLU1 catalytic domains, plus a mannosyltransferase domain and a pyridine nucleotide disulfide oxidoreductase domain in the N-terminal and C-terminal regions, respectively. This NADH-GOGAT was constitutively transcribed at an approximate 2.5-fold lower level than the g70690 encoded Fd-GOGAT (Fig. 4G). The g206190 encoded NADH-GOGAT was not expressed. Thus, there are one each Fd- and NADH-dependent GOGAT proteins constitutively present in *A*. *punctatus* under our experimental conditions. In tracheophytes, enhanced transcription of GOGAT encoding genes is tissue specific in response to photosynthate and N metabolites (Suzuki and Knaff, 2005) The genes encoding Fd-GOGAT are upregulated by light (especially red), sucrose, nitrate, ammonium, and other substrates and products of the GS-GOGAT pathway; conversely the NADH-GOGAT encoding genes appears to be primarily under nitrate and ammonium control with the most robust expression in roots. (Suzuki and Knaff, 2005).

Collectively, these data provide support for N-control over key enzymes of N-assimilation; specifically ammonium and nitrate transport, and a plastid GS2 plus a moderately expressed constitutive nodGS (encoded by g14110) with a signal peptide that implies compartmentalization. Moreover, they illustrate that the complexity of N-assimilation in bryophytes is like that in seed plants.

#### Potential sinks for excess reductant accumulation

Candidates for organic end products that function as internally stored or excreted reductant sinks during N-starvation include products such as polysaccharides, fatty acids and lipids, and secondary metabolites. We predict expression patterns for such sinks should follow those shown in cluster 1, where transient upregulation of gene expression was complemented in correlation with the onset of symbiotic N_2_ fixation.

Since the symbiotic cavity in gametophyte tissue contains considerable slime, secretory polysaccharide synthesis must happen, and upregulation of the pathway would be a relatively benign solution to the consequences of N-starvation. Neither the composition nor the biosynthetic pathway of cavity slime is not known; nevertheless, the differential transcription patterns of the glycosyltransferases in *A. punctatus* are not consistent with a role in increased slime production (an example is shown in Fig. 5A). Triacylglycerol (TAG) and lipid accumulation are well documented in microalgae during N-starvation (Goncalves *et al.*, 2016) as well as in seeds of certain plants, such as castor bean (Chen *et al.*, 2018*a*). The genome contains 14 genes encoding acyl carrier proteins involved in both fatty acid and TAG synthesis; one, identified as encoding Kas1, is present in the differential transcriptome in cluster 3; two were constitutively transcribed and the remainder were not expressed. Relative to other plants and algae, fatty acid, TAG and lipid accumulation do not appear to be robust during N-starvation of *A. punctatus*. These possibilities will need to be addressed biochemically.

**Figure 5:**
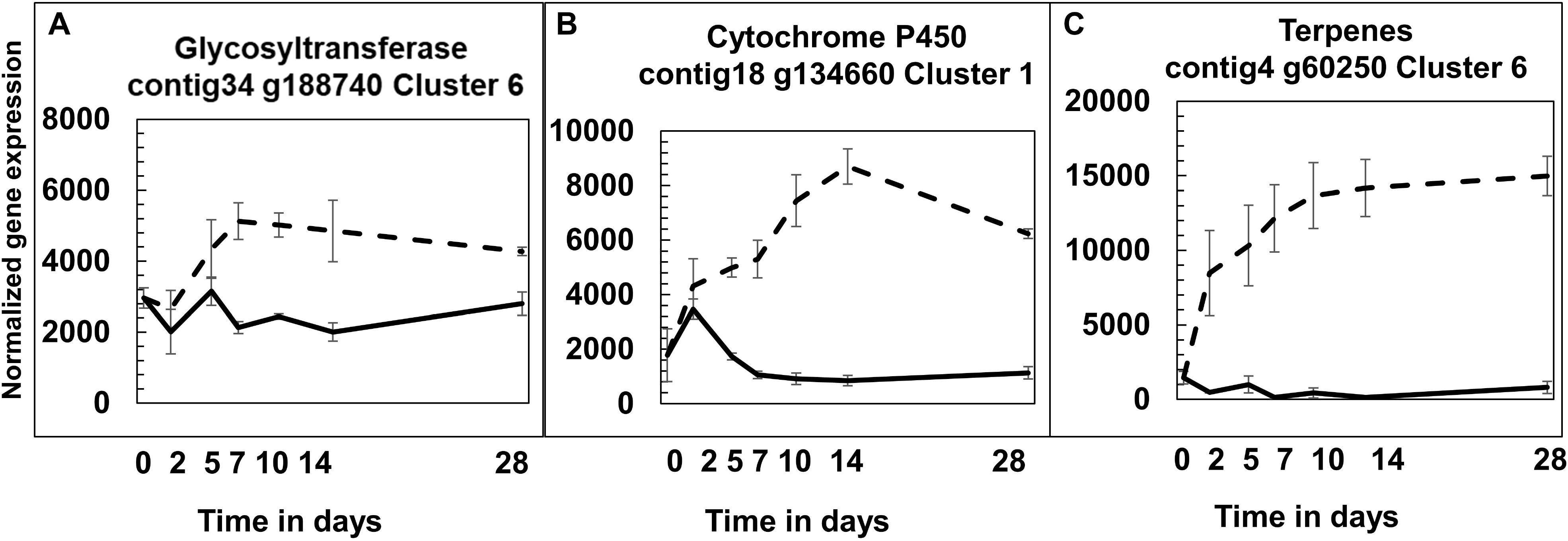
Temporal patterns of normalized expression for genes involved in potential reductant sinks for photosynthetically generated reductant. The ordinate is the mean (±SE) normalized gene expression of transcripts (P<0.05). The selected genes encode: glycosyltransferase (A), a cytochrome P450 (B), terpenes (C), from cocultured and symbiont-free gametophytes. Time is in days as in Fig. 2.

Genes encoding cytochrome P450 (CYP450) were abundant in the differential transcriptome and highly represented in clusters 1 and 6. Only those in cluster 1 would be consistent with a transient N-starvation response complemented by N_2_-derived ammonium. CYP450 is a monooxygenase whose activities are involved in numerous metabolic pathways, especially those leading to stress adaptation, development, and synthesis of secondary metabolites (Xu *et al*., 2015). An example of the temporal expression pattern of one CYP450 is presented in Fig. 5B, where it shows initial upregulation in the absence and presence of *N. punctiforme*. The observation that the presence of *N. punctiforme* negated the upregulation, in a pattern approximating that of induction of symbiotic nitrogenase activity, is consistent with an enzyme that is specific for N-stress in synthesis of an electron sink. It should be noted that at relatively larger number of CYP450 encoding genes are also differentially transcribed in the pattern of cluster 6 (e.g. g57450), where the presence of *N*. *punctiforme* repressed transcription before nitrogenase activity had been induced (Dataset S3). These results indicate that specific CYP450 proteins are highly present under N-stress. Terpenes are a fundamental substrate for synthesis of a variety of stable and volatile secondary plant products that could well function as electron sinks. Moreover, the products are synthesized following a variety of abiotic and biotic stresses (Chatterjee *et al*., 2017; 2018; 2020). One example of the differential expression of a terpene synthase from cluster 6 in shown in Fig. 5C; the presence of *N. punctiforme* completely suppressed upregulation of transcription from the marginal time zero level. Glycosylation is also involved in the biosynthesis and storage of secondary compounds. In plants, these reactions are controlled by a specific subclass of the ubiquitous glycosyltransferase family (Tiwari *et al.*, 2016). The upregulated transcription pattern in symbiont-free *A. punctatus* is consistent with a role of glycosylation under N-starvation, whereas cocultured *A. punctatus* ameliorated the stress response (Fig. 4F). However, we provide no evidence for the substrate of this glycosyltransferase.

### Genes Related to Symbiotic Interactions

Based on morphological, physiological and biochemical data, we previously suggested that the signaling between hornwort and cyanobacterium is primarily unidirectional from plant to symbiont, excluding transfer of N_2_-derived ammonium (Meeks, 1998). The temporal patterns of the diametrically opposite clusters 2 and 6 contradict that suggestion. The upregulation of DEGs in cluster 2 and their downregulation in cluster 6 were apparent by day 2 of coculture, during which time *N*. *punctiforme* hormogonia were colonizing the slime cavity, prior to the provision of fixed N, which was argued to contribute to the temporal patterns in clusters 1, 3 and 7. We do not yet know whether the early signaling is chemical or physical contact but have experiments to clarify this in progress using existing *N*. *punctiforme* mutants that differentiate hormogonia which are unable to infect gametophyte tissue and mutants that are infective but unable to fix N_2_.

A central question regarding the hornwort-cyanobacteria N_2_-fixing symbiosis is whether proteins constituting the plant CSSP are involved in establishment of the association. Nine genes encoding proteins of the CSSP (Oldroyd, 2013; Sellstedt and Richau, 2013; Debellé, 2020; Delaux and Schornack, 2021) were reported present in the *A. punctatus* genome (Li *et al.*, 2020). We found none of those genes in our differentially expressed transcriptome. We searched the transcriptome (Dataset S1) to determine if they were constitutively transcribed. This led to the detection of the nine CSSP encoding genes (Fig. 5A-J): Castor (calcium channel), Cyclops (calcium calmodulin-dependent binding to CCamK), STR1 (ABC transporter), STR2 (ABC transporter), SymRK (receptor-mediated signaling), CCamK (calcium and calmodulin-dependent serine/threonine protein kinase), Vapyrin (protein kinase), RAD1 (GRAS family transcription factor [TF]) and one copy of RAM1 (GRAS family TF) were expressed (Fig. 6A-I). RAM1 displays a temporal pattern of cluster 1 but was not sorted as differentially expressed. RAD1 was expressed at a relatively low level, and a second copy of RAM1 was not expressed. A LysM receptor kinase is involved in perception of nodulation factors (Buendia *et al.*, 2018) and a gene putatively encoding such a protein was present in the transcriptome at a substantially high expressed level (Fig. 5J). These results indicate that the sensing and immediate signaling capacities of the CSSP are present in gametophyte tissue to recognize a compatible symbiont, which may or may not be *Nostoc* spp. Since hornworts do form mycorrhizal associations (Desirò *et al.*, 2013), we suggest the CSSP in *A. punctatus* may be restricted to its originally evolved role (Oldroyd, 2013). This is consistent with our previous suggestion that each group of symbiotic land plant partners of N_2_-fixing cyanobacteria (overwhelmingly *Nostoc* spp.) evolved different mechanisms to achieve control over the same metabolic processes in the cyanobacterium while arriving at a stable, competitive symbiotic association (Meeks, 1998). Nevertheless, because the genes are constitutively transcribed at a high level, involvement of the CSSP in cyanobacterial symbiosis with hornworts needs to be studied by mutational analysis, which is now being developed in *A*. *agrestis* (Frangedakis *et al.*, 2021).

**Figure 6:**
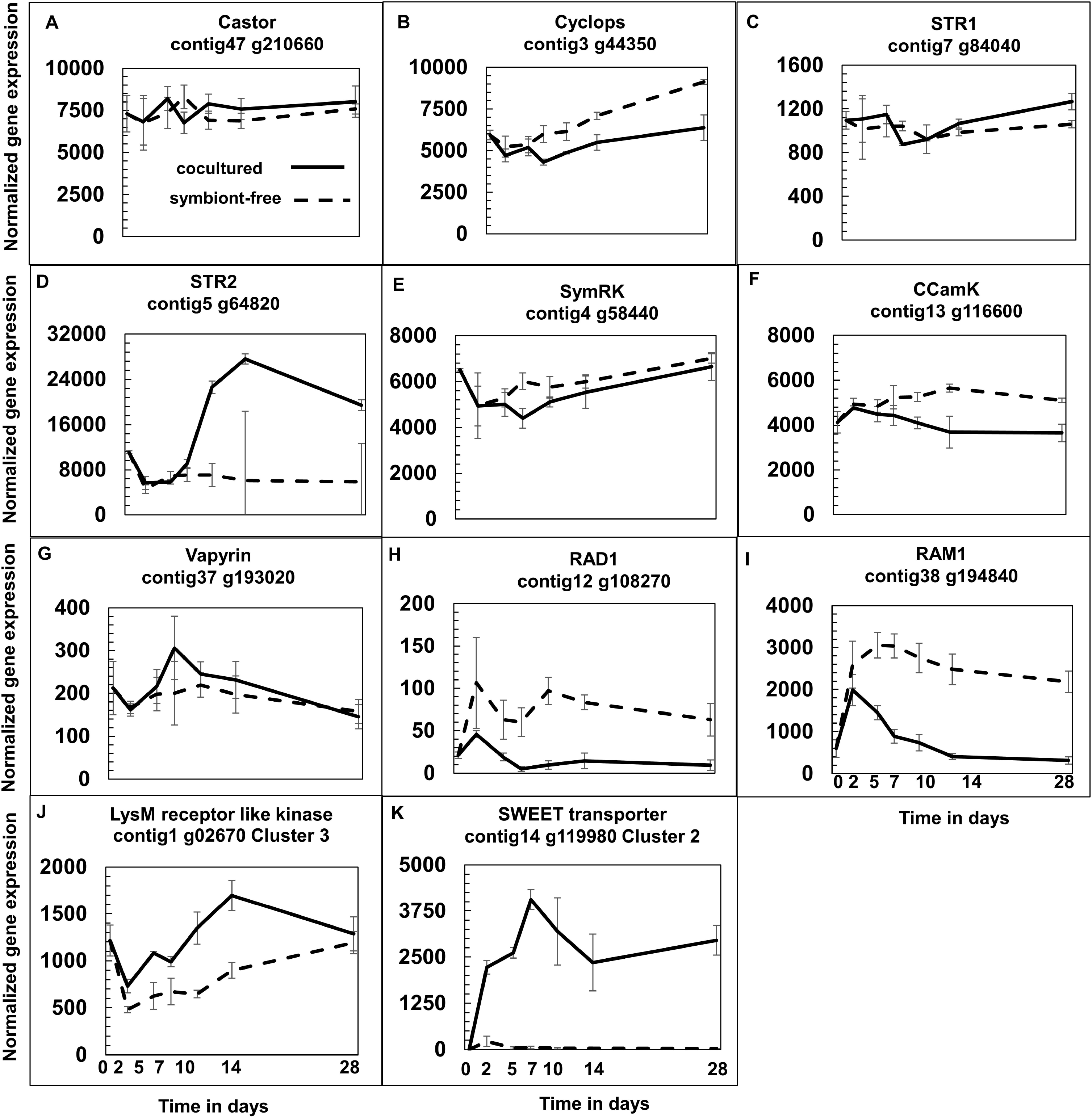
Temporal patterns of normalized expression for genes involved in symbiotic interactions. The ordinate is mean (±SE) of normalized expression of genes encoding the proteins: Castor (A), Cyclops (B), STR1 (C), STR2 (D), SymRK (E), CCamK (F), Vapyrin (G), RAD1 (H), RAM1 (I), LysM receptor like kinase (J), SWEET transporter (K) from cocultured and symbiont-free *A. punctatus* gametophytes. Time is in days as in Fig. 2. All data except LysM (J) and SWEET (K) were obtained from the transcriptome. LysM and SWEET sorted into the differential transcriptome, while STR2 and RAM1 did not.

Transcription of the gene for a sugar transporter (SWEET) in *A. punctatus* was previously identified as specifically upregulated in the presence of *N*. *punctiforme* (Li *et al.*, 2020) and verified here (Fig. 6K). What extends the prior observation is the fact that the activation of transcription of SWEET occurred early in the presence of *N*. *punctiforme*, prior to its provision of N_2_-derived ammonium, implying transcription may not initially be N-starvation dependent.

The implication of *N*. *punctiforme* dependent upregulation was the hypothesis that *A*. *punctatus* has a role in providing sugars, such as sucrose, fructose and glucose that are known to support symbiotic N_2_-fixation by *Nostoc* spp. in association with *A*. *punctatus* (Steinberg and Meeks, 1991). In addition, we have observed a dependence on a *N*. *punctiforme* glucose permease for development of a functional symbiotic association (Ekman *et al.*, 2013).

To summarize, with our much-improved genome assembly and detailed temporal RNA-seq experiments, we found evidence of bidirectional communications between the partners and, for the first time, revealed how symbiotic cyanobacteria impact plant hosts’ global gene expression through time. We also extend to hornworts knowledge of the complexity of N acquisition and assimilation. With the recent development of a hornwort transformation system, it is becoming feasible to carry out gene functional studies. We anticipate the candidate genes, and their putative functions, uncovered here will form the necessary foundation for future investigations into the genetics of cyanobacteria symbiosis.

## Supporting information

Supplementary Dataset S1-S3+ Supplementary Table S1+ Supplementary guide S1+ Figures S1-S2

## Abbreviations

CSSP: common symbiotic signaling pathway
HIF: hormogonium inducing factor

## Supplementary Data

**Supplementary Guide S1:** Guide to analysis of functional assignments of differentially expressed genes.

**Supplementary Figure 1**. PCA analysis of biological replicates.

Control (T0), cocultured (T2IN, T5IN, T7IN, T10IN, T14IN and T28IN) and symbiont-free (T2UN, T5UN, T7UN, T10UN, T14UN and T28UN) *A. punctatus* under N-starved conditions during time course RNA-Seq analysis.

**Supplementary Figure 2.** Manually assigned metabolic categories of differentially transcribed genes observed in cluster analysis.

**Supplementary Table 1.** Length, expression and domain organization of multiple gene families involved in N acquisition and assimilation.

**Supplementary Dataset S1**. Gene count matrix containing all the transcript counts of 23,021 genes from 39 samples.

**Supplementary Dataset S2**. The functional annotation file containing all KEGG and GO annotations for 69% of the transcripts from gene count matrix.

**Supplementary Dataset S3**. The list of 1,210 significantly differentially expressed genes as an output from the cluster analysis, as well as the clustered distribution of each gene and its classification into ten major metabolic categories.

## Acknowledgements

This research was supported by the USA National Science Foundation grant no. DEB1831552 to JCM and DEB1831428 to FWL. We thank Steven M. Theg and Matthew Edmund Gilbert for loan of the PAM fluorometer and Miguel A. Valderrama Gomez for assistance in computational analyses. We thank Janet Sprent for the inspiration of “*Nostoc* talks back”.

## Author contribution

PC and JCM: design of the research; PC: performance of the research; PC, FWL, PS: data analysis; PC, JCM: interpretation; PC and JCM: writing the manuscript, with input from FWL and PS.

## Data availability

All primary data for the RNA sequencing are deposited at SRA NCBI Genbank with accession no. PRJNA750572

## References

Adams DG. 2002. Cyanobacteria in symbiosis with hornworts and liverworts. In Cyanobacteria in symbiosis (pp. 117–135). Springer, Dordrecht.

Brůna T, Hoff KJ, Lomsadze A, Stanke M, Borodovsky M. 2021. BRAKER2: automatic eukaryotic genome annotation with GeneMark-EP+ and AUGUSTUS supported by a protein database. NAR Genomics and Bioinformatics 3.

Buendia L, Girardin A, Wang T, Cottret L, Lefebvre B. 2018. LysM receptor-like kinase and LysM receptor-like protein families: an update on phylogeny and functional characterization. Frontiers in Plant Science. 9, 1531.

Calabrese S, Kohler A, Niehl A, Veneault-Fourrey C, Boller T, Courty P-E. 2017. Transcriptome analysis of the P*opulus trichocarpa*–*Rhizophagus irregularis* Mycorrhizal Symbiosis: Regulation of plant and fungal transportomes under nitrogen starvation. Plant and Cell Physiology 58, 1003–1017.

Campbell EL, Meeks JC. 1989. Characteristics of hormogonia formation by symbiotic *Nostoc* spp. in response to the presence of *Anthoceros punctatus* or its extracellular products. Applied and Environmental Microbiology 55, 125–131.

Cantalapiedra CP, Hernández-Plaza A, Letunic I, Bork P, Huerta-Cepas J. 2021. eggNOG-mapper v2: Functional annotation, orthology assignments, and domain prediction at the metagenomic scale. BioRxiv, 2021.06.03.446934.

Chatterjee P, Kanagendran A, Samaddar S, Pazouki L, Sa TM, Niinemets Ü. 2020. Influence of *Brevibacterium linens* RS16 on foliage photosynthetic and volatile emission characteristics upon heat stress in *Eucalyptus grandis*. Science of The Total Environment 700, 134453.

Chatterjee P, Kanagendran A, Samaddar S, Pazouki L, Sa TM, Niinemets Ü. 2018. Inoculation of *Brevibacterium linens* RS16 in *Oryza sativa* genotypes enhanced salinity resistance: impacts on photosynthetic traits and foliar volatile emissions. Science of the Total Environment 645, 721–732.

Chatterjee P, Samaddar S, Anandham R, Kang Y, Kim K, Selvakumar G, Sa T. 2017. Beneficial soil bacterium *Pseudomonas frederiksbergensis* OS261 augments salt tolerance and promotes red pepper plant growth. Frontiers in plant science, 8, 705.

Chellamuthu VR, Alva V, Forchhammer K. 2013. From cyanobacteria to plants: conservation of PII functions during plastid evolution. Planta 237, 451–462.

Chen Y, Cen K, Lu Y, Zhang S, Shang Y, Wang C. 2018a. Nitrogen-starvation triggers cellular accumulation of triacylglycerol in *Metarhizium robertsii*. Fungal Biology 122, 410–419.

Chen S, Zhou Y, Chen Y, Gu J. 2018b. fastp: an ultra-fast all-in-one FASTQ preprocessor. Bioinformatics 34, i884–i890.

Cohen MF, Wallis JG, Campbell EL, Meeks JC. 1994. Transposon mutagenesis of *Nostoc* sp. strain ATCC 29133, a filamentous cyanobacterium with multiple cellular differentiation alternatives. Microbiology 140, 3233–3240.

Cohen MF, Meeks JC. 1997. A hormogonium regulating locus, hrmUA, of the cyanobacterium *Nostoc punctiforme* strain ATCC 29133 and its response to an extract of a symbiotic plant partner *Anthoceros punctatus*. Molecular plant-microbe interactions: MPMI 10, 280–289.

Cohen MF, Meeks JC, Cai YA, Wolk CP. 1998. Transposon mutagenesis of heterocyst-forming filamentous cyanobacteria. Methods in Enzymology. Academic Press, 3–17.

Conesa A, Nueda MJ, Ferrer A, Talón M. 2006. maSigPro: a method to identify significantly differential expression profiles in time-course microarray experiments. Bioinformatics 22, 1096–1102.

Couturier J, Montanini B, Martin F, Brun A, Blaudez D, Chalot M. 2007. The expanded family of ammonium transporters in the perennial poplar plant. The New Phytologist 174, 137–150.

Crawford NM, Glass ADM. 1998. Molecular and physiological aspects of nitrate uptake in plants. Trends in Plant Science 3, 389–395.

Debellé F. 2020. The common symbiotic signaling pathway. The model legume Medicago truncatula. John Wiley & Sons, Ltd, 523–528.

Delaux P-M, Schornack S. 2021. Plant evolution driven by interactions with symbiotic and pathogenic microbes. Science 371.

Desirò A, Duckett JG, Pressel S, Villarreal JC, Bidartondo MI. 2013. Fungal symbioses in hornworts: a chequered history. Proceedings Of The Royal Society B: Biological Sciences 280, 20130207.

Ekman M, Picossi S, Campbell EL, Meeks JC, Flores E. 2013. A *Nostoc punctiforme* sugar transporter necessary to establish a Cyanobacterium-Plant symbiosis. Plant Physiology 161, 1984–1992.

Enderlin CS, Meeks JC. 1983. Pure culture and reconstitution of the *Anthoceros-Nostoc* symbiotic association. Planta 158, 157–165.

Feller U, Anders I, Mae T. 2008. Rubiscolytics: fate of Rubisco after its enzymatic function in a cell is terminated. Journal of Experimental Botany 59, 1615–1624.

Frangedakis E, Waller M, Nishiyama T, et al. 2021. An *Agrobacterium*-mediated stable transformation technique for the hornwort model *Anthoceros agrestis*. New Phytologist

Ghodhbane-Gtari F, Hurst SG, Oshone R, Morris K, Abebe-Akele F, Thomas WK, Ktari A, Salem K, Gtari M, Tisa LS. 2014. Draft Genome Sequence of *Frankia* sp. Strain BMG5.23, a Salt-Tolerant nitrogen-fixing Actinobacterium isolated from the root nodules of *Casuarina glauca* grown in Tunisia. Genome Announcements 2.

Glass ADM. 2002. The regulation of nitrate and ammonium transport systems in plants. Journal of Experimental Botany 53, 855–864.

Goncalves EC, Wilkie AC, Kirst M, Rathinasabapathi B. 2016. Metabolic regulation of triacylglycerol accumulation in the green algae: identification of potential targets for engineering to improve oil yield. Plant Biotechnology Journal 14, 1649–1660.

Guo J, Jia Y, Chen H, Zhang L, Yang J, Zhang J, Hu X, Ye X, Li Y, Zhou Y. 2019. Growth, photosynthesis, and nutrient uptake in wheat are affected by differences in nitrogen levels and forms and potassium supply. Scientific Reports 9, 1248.

Harris JM, Pawlowski K, Mathesius U. 2020. Editorial: Evolution of Signaling in Plant Symbioses. Frontiers in Plant Science 11, 456.

Horváth B, Yeun LH, Domonkos Á, et al. 2011. *Medicago truncatula* IPD3 is a member of the common symbiotic signaling pathway required for Rhizobial and Mycorrhizal symbioses. Molecular Plant-Microbe Interactions 24, 1345–1358.

Hsieh MH, Lam HM, Van De Loo FJ, Coruzzi G. 1998. A PII-like protein in *Arabidopsis*: putative role in nitrogen sensing. Proceedings of the National Academy of Sciences 95, 13965–13970.

Hsieh P-H, Kan C-C, Wu H-Y, Yang H-C, Hsieh M-H. 2018. Early molecular events associated with nitrogen deficiency in rice seedling roots. Scientific Reports 8, 12207.

Inskeep WP, Bloom PR. 1985. Extinction coefficients of chlorophyll *a* and *b* in *N,N* - Dimethylformamide and 80% Acetone. Plant Physiology 77, 483–485.

Kim D, Langmead B, Salzberg SL. 2015. HISAT: a fast spliced aligner with low memory requirements. Nature methods 12, 357–360.

Kolmogorov M, Yuan J, Lin Y, Pevzner PA. 2019. Assembly of long, error-prone reads using repeat graphs. Nature Biotechnology 37, 540–546.

Krapp A, Berthomé R, Orsel M, Mercey-Boutet S, Yu A, Castaings L, Elftieh S, Major H, Renou J-P, Daniel-Vedele F. 2011. Arabidopsis roots and shoots show distinct temporal adaptation patterns toward nitrogen starvation. Plant Physiology 157, 1255–1282.

Latowski D, Kuczyńska P, Strzałka K. 2011. Xanthophyll cycle--a mechanism protecting plants against oxidative stress. Redox Report: Communications in Free Radical Research 16, 78–90.

Li F-W, Nishiyama T, Waller M, et al. 2020. *Anthoceros* genomes illuminate the origin of land plants and the unique biology of hornworts. Nature Plants 6, 259–272.

Liu W, Sun Q, Wang K, Du Q, Li W-X. 2017. Nitrogen limitation adaptation (NLA) is involved in source-to-sink remobilization of nitrate by mediating the degradation of NRT1.7 in *Arabidopsis*. New Phytologist 214, 734–744.

Logan BA, Demmig-Adams B, Rosenstiel TN, AdamsIII WW. 1999. Effect of nitrogen limitation on foliar antioxidants in relationship to other metabolic characteristics. Planta, 209(2), pp.213–220.

Loqué D, von Wirén N. 2004. Regulatory levels for the transport of ammonium in plant roots. Journal of Experimental Botany 55, 1293–1305.

de Marsac NT. 1994. Differentiation of hormogonia and relationships with other biological processes. In: Bryant DA, ed. Advances in Photosynthesis. The Molecular Biology of Cyanobacteria. Dordrecht: Springer Netherlands, 825–842.

McDonald TR, Ward JM. 2016. Evolution of electrogenic ammonium transporters (AMTs). Frontiers in Plant Science 7, 352.

Meeks JC. 1998. Symbiosis between nitrogen-fixing cyanobacteria and plants. BioScience 48, 266–276.

Meeks JC, Castenholz RW. 1971. Growth and photosynthesis in an extreme thermophile,*Synechococcus lividus* (Cyanophyta). Archiv für Mikrobiologie 78, 25–41.

Meeks JC, Enderlin CS, Joseph CM, Chapman JS, Lollar MWL. 1985. Fixation of [13N]N_2_ and transfer of fixed nitrogen in the *Anthoceros-Nostoc* symbiotic association. Planta 164, 406–414.

Meeks JC, Enderlin CS, Wycoff KL, Chapman JS, Joseph CM. 1983. Assimilation of 13NH_4_ ^+^by *Anthoceros* grown with and without symbiotic *Nostoc*. Planta 158, 384–391.

Meeks JC. 2003. Symbiotic interactions between *Nostoc punctiforme*, a multicellular cyanobacterium, and the hornwort *Anthoceros punctatus*. Symbiosis (Philadelphia, PA), 35, 55–71.

Mertz IT, Christians NE, Thoms AW. 2019. Branched-chain amino acids for use as a nitrogen source on creeping Bentgrass. HortTechnology 29, 833–837.

Miller AJ, Fan X, Shen Q, Smith SJ. 2008. Amino acids and nitrate as signals for the regulation of nitrogen acquisition. Journal of Experimental Botany 59, 111–119.

Murchie EH, Lawson T. 2013. Chlorophyll fluorescence analysis: a guide to good practice and understanding some new applications. Journal of Experimental Botany 64, 3983–3998.

Mus F, Crook MB, Garcia K, et al. 2016. Symbiotic nitrogen fixation and the challenges to its extension to nonlegumes. Applied and Environmental Microbiology 82, 3698–3710.

Nilsson M, Rasmussen U, Bergman B. 2006. Cyanobacterial chemotaxis to extracts of host and nonhost plants. FEMS microbiology ecology, 55(3), 382–390.

Niyogi KK, Björkman O, Grossman AR. 1997. The roles of specific xanthophylls in photoprotection. Proceedings of the National Academy of Sciences of the United States of America 94, 14162–14167.

O’Brian MR. 1996. Heme synthesis in the rhizobium-legume symbiosis: a palette for bacterial and eukaryotic pigments. Journal of Bacteriology 178, 2471–2478.

Oldroyd GED. 2013. Speak, friend, and enter: signalling systems that promote beneficial symbiotic associations in plants. Nature Reviews Microbiology 11, 252–263.

Ou S, Su W, Liao Y, et al. 2019. Benchmarking transposable element annotation methods for creation of a streamlined, comprehensive pipeline. Genome Biology 20, 275.

Pankievicz VCS, Irving TB, Maia LGS, Ané J-M. 2019. Are we there yet? The long walk towards the development of efficient symbiotic associations between nitrogen-fixing bacteria and non-leguminous crops. BMC Biology 17, 99.

Paul MJ, Driscoll SP. 1997. Sugar repression of photosynthesis: the role of carbohydrates in signalling nitrogen deficiency through source:sink imbalance. Plant, Cell & Environment 20, 110–116.

Paungfoo-Lonhienne C, Lonhienne TGA, Rentsch D, Robinson N, Christie M, Webb RI, Gamage HK, Carroll BJ, Schenk PM, Schmidt S. 2008. Plants can use protein as a nitrogen source without assistance from other organisms. Proceedings of the National Academy of Sciences 105, 4524–4529.

Pérez-Jaramillo JE, Mendes R, Raaijmakers JM. 2016. Impact of plant domestication on rhizosphere microbiome assembly and functions. Plant Molecular Biology 90, 635–644.

Pertea M, Pertea GM, Antonescu CM, Chang T-C, Mendell JT, Salzberg SL. 2015. StringTie enables improved reconstruction of a transcriptome from RNA-seq reads. Nature Biotechnology 33, 290–295.

Rodgers GA, Stewart WDP. 1977. The Cyanophyte-Hepatic Symbiosis I. Morphology and Physiology. New Phytologist 78, 441–458.

Sayed OH. 1998. Analysis of photosynthetic responses and adaptation to nitrogen starvation in Chlorella using in vivo chlorophyll fluorescence. Photosynthetica 35, 611–619.

Sellstedt A, Richau KH. 2013. Aspects of nitrogen-fixing Actinobacteria, in particular free-living and symbiotic Frankia. FEMS Microbiology Letters 342, 179–186.

Simão FA, Waterhouse RM, Ioannidis P, Kriventseva EV, Zdobnov EM. 2015. BUSCO: assessing genome assembly and annotation completeness with single-copy orthologs. Bioinformatics 31, 3210–3212.

Smit AFA, Hubley R, Green P. 2013-2015. RepeatMasker Open-4.0.

Sprent JI, Raven JA. 1985. Evolution of nitrogen-fixing symbioses. Proceedings of the Royal Society of Edinburgh, Section B: Biological Sciences 85, 215–237.

Steinberg NA, Meeks JC. 1991. Physiological sources of reductant for nitrogen fixation activity in *Nostoc* sp. strain UCD 7801 in symbiotic association with *Anthoceros punctatus*. Journal of Bacteriology 173, 7324–7329.

Sun J, Zheng N. 2015. Molecular mechanism underlying the plant NRT1.1 dual-affinity nitrate transporter. Frontiers in Physiology 6.

Suzuki A, Knaff DB. 2005. Glutamate synthase: structural, mechanistic and regulatory properties, and role in the amino acid metabolism. Photosynthesis Research 83, 191–217.

Swarbreck SM, Defoin-Platel M, Hindle M, Saqi M, Habash DZ. 2011. New perspectives on glutamine synthetase in grasses. Journal of Experimental Botany 62, 1511–1522.

Szövényi P, Gunadi A, Li FW. 2021. Charting the genomic landscape of seed-free plants. Nature Plants 7, 554–565.

Tiwari P, Sangwan RS, Sangwan NS. 2016. Plant secondary metabolism linked glycosyltransferases: An update on expanding knowledge and scopes. Biotechnology Advances 34, 714–739.

Walker BJ, Abeel T, Shea T, et al. 2014. Pilon: an integrated tool for comprehensive microbial variant detection and genome assembly improvement. PloS One 9, e112963.

Walsby AE. 2007. Cyanobacterial heterocysts: terminal pores proposed as sites of gas exchange. Trends in Microbiology 15, 340–349.

Wang YY, Cheng YH, Chen KE, Tsay YF. 2018. Nitrate Transport, Signaling, and Use Efficiency. Annual Review of Plant Biology 69, 85–122.

Jun XU, Wang XY, Guo WZ. 2015. The cytochrome P450 superfamily: Key players in plant development and defense. Journal of Integrative Agriculture 14, 1673–1686.

Yandell B. 2017. Practical Data Analysis for Designed Experiments. Routledge.

Zalutskaya Z, Kochemasova L, Ermilova E. 2018. Dual positive and negative control of *Chlamydomonas* PII signal transduction protein expression by nitrate/nitrite and NO via the components of nitric oxide cycle. BMC Plant Biology 18, 305.

Zhao L-S, Li K, Wang Q-M, et al. 2017. Nitrogen starvation impacts the photosynthetic performance of P*orphyridium cruentum* as revealed by chlorophyll a fluorescence. Scientific Reports 7, 8542.

