## Supplementary Dataset S1-S3+ Supplementary Table S1+ Supplementary guide S1+ Figures S1-S2 for "*Nostoc* talks back: Temporal patterns of differential gene expression during establishment of the *Anthoceros-Nostoc* symbiosis": Supplementary Table S1+ Supplementary guide S1+ Figures S1-S2.pdf

**Supplementary Table S1.** Length, expression and domain organization of multiple gene families involved in N acquisition and assimilation.

| Gene location number Anthoceros_punctatus_v2 | Gene symbol | Amino acids | Cluster No/ Expression level | Protein domains |
|---|---|---|---|---|
| Ammonium | | | | |
| Contig 3_g54810 | AMT1;5 | 316 | 1/39836* | 45-312 Ammonium/urea transport; AMTB-like |
| Contig27_g162610 | AMT2;1 | 575 | Con/28121** | 23-457 Ammonium transport; AMTB-like; 145-344 Rhesus RHD |
| Contig34_g187510 | AMTB | 482 | Con/482** | 15-424 Ammonium transport; AMTB-like; 118-353 Rhesus RHD |
| Contig54_g215080 | fusion | 1774 | Con/3862** | 30-459 and 930-1327 Ammonium transport; 522-821 and 1409-1708 PP2C-type Phosphatase |
| Contig35_gG189910 | AMTB | 480 | Con/1.05** | 34-462 Ammonium/urea transport; AMTB-like |
| Contig66_g219420 | AMTB | 420 | Not expressed | 9-410 Ammonium/urea transporter; AMTB-like |
| Contig70_g221650 | AMTB | 420 | Not expressed | 6-410 Ammonium transport; AMTB-like; 10-315 Rhesus RHD |
| Nitrate | | | | |
| Contig14_g119600 | NRT2 | 356 | Con/23382 | 15-530 High affinity nitrate transporter 2;3 or 2;4 (MFS) |
| Contig16_g128250 | NRT2 fragment | 188 | Com/38702 | 8-184 High affinity nitrate transporter 3.2; 1-23 signal peptide; no transmembrane domains in transporter domain |
| Glutamine synthetase | | | | |
| Contig25_g158060 | GS2 | 413 | 3/3124* | 10-442 Glutamine synthetase; 2-98 Gln synth N-terminal; 99-439 Gln synth catalytic domain; probable signal peptide |
| Contig16_g128160 | GS1;1 | 315 | Con/61342** | 1-315 Glutamine synthetase; 22-96 Gln synth b-grasp; 102-315 Gln synth catalytic domain |
| Contig19_g141120 | nodGS | 532 | Con/1552** | 77-530 Glutamine synthetase; 77-190 Gln synth N-terminal; 191-528 Gln synth catalytic domain; 1-32 signal peptide |
| Contig3_g45770 | nodGS | 443 | Con/466** | 10-442 Glutamine synthetase; 2-98 Gln synth N-terminal; 99-439 Gln synth catalytic domain; 1-22 signal peptide; GS type-III |
| Contig11_g103610 | fusion | 819 | Con/3748** | 41-405 Metallohydrolase; 186-386 Amidohydrolase; 483-810 Gln synth catalytic/guanido kinase; 1-37 signal peptidase. |
| Contig11_g105000 | nodGS | 131 | Con/27 | 1-31 N-terminal FLUG-like |
| Contig 65_g219110 | GS1 | 437 | Not expressed | 3-436 glutamine synthetase; 16-97 Gln synth b-grasp; 104-434 Gln synth catalytic domain |
| Contig65_g219070 | GS1 | 445 | Not expressed | 8-440 glutamine synthetase; 17-98 Gln synth b-grasp; 104-439 Gln synth catalytic domain. |
| Glutamate synthase | | | | |

| | | | | |
|---|---|---|---|---|
| Contig6_g70690 | Fd-GOGAT | 1265 | Con/17995** | 1-1265 Glutamate synthase; 1-171 GATase domain; 198-490 Glu Synth central domain; 550-948 Glu Synth; 990-1240 Glu Synth C-terminal |
| Contig6_g70700 | Fd-GOGAT fragment | 329 | Con/1558** | 1-329 Glutamate synthase; 1-21 signal peptide; 138-329 GATase type 2 domain |
| Cintig5_g63560 | NADH-GOGAT | 2462 | Con/6464** | 314-2002 Glutamate synthase 1; 1-314 mannosyltransferase (TM domains); 350-778 GATase; 823-1111 Glu Synth central N; 1180-1547 Glu synth; 1608-1856 Glu synth (alpha subunit); 2003-2379 NAD binding |
| Contig41_g206190 | NADH-GOGAT | 2277 | Not expressed | 299-2277 Glutamate synthase; 20-181 glycosyltransferase (TM domains); 288-398 GTRA; 469-882 GATase; 919-1199 Glu synth central N; 1474-1777 NAD(P) binding; 1891-2059 Glu synth (alpha subunit) |

Expression level is *average of normalized counts from symbiotic tissue; **average of normalized counts from symbiont-free tissue. Con is constitutive expression. AMTB-like identifies prokaryotic associated genes. Fusion is presence of domains not associated with assigned activity. MFS is major facilitator superfamily. TM is transmembrane domains.

**Supplementary Guide S1**

Guide to the manually organized differential transcriptome

Unassigned functions contain the highest category with 390 transcripts. These transcripts may have been identified by a known domain but unidentified function, have a homolog of unknown function observed in other organisms, or be recognized as encoding a hypothetical protein. This category represents 27% of the differentially expressed transcriptome.

Core metabolism (permissive steady state synthesis of precursors, monomers, polysaccharides, lipids and steroids, oxidation/reduction reactions, secondary metabolites and nutrient reservoirs, amongst others) constitutes the next most highly represented category with 360 different transcripts. The 67 secondary metabolite encoding genes were the highest of the subgroups but 72% of these transcripts encode cytochrome P450. While present in all clusters, except for 2 and 8, CYP450 transcripts predominated in clusters 1 (18 transcripts) and 6 (20 transcripts); these two clusters showed upregulation and persistence of expression in the absence of *N. punctiforme*, consistent with gene products important for survival under the stressed conditions. Other more highly enriched core metabolism subcategories include those encoding oxidation/reduction reactions (42 transcripts, with 52% in clusters 1 and 6); precursor carbon metabolic pathways (41 transcripts with 1 to 8 different transcripts in all clusters but 8); fatty acid and lipid synthesis (41 transcripts with 24% in cluster 7); glycosyltransferases (26 transcripts with 1 to 6 transcripts in all clusters except 8 and 9); amino acids (21 transcripts with 57% in clusters 2 and 3 and the remainder distributed in the other clusters except for none in 1, 8 and 9); and nutrient storage (16 transcripts with 75% in clusters 5 and 7, none in 6, 8 and 9, and single transcripts in the remaining clusters).

The three categories of cellular processes (122 transcripts), stress, defense and detoxification (121 transcripts) and protein metabolism (119 transcripts) each account for approximately 10% of the total differentially expressed transcripts (1,210). The cellular processes subgroups include, amongst others: **i)** Chromosome biology, nuclear activities, and DNA replication and repair of which 45 transcripts are present in all clusters except 8, but the transcripts are enriched in clusters 4 and 9 with 8 and 20 transcripts, respectively; clusters 4 and 9 showed similar early burst and decline patterns of transcription in the absence and presence of *N. punctiforme*. **ii)** Cell walls with

25 transcripts in all clusters except 4 and 9 and 52% in clusters 1, 3 and 7, which display two different temporal patterns. **iii)** Cell trafficking with 22 transcripts were distributed relatively evenly in all clusters except 5, 6 and 8.

Stress (19 transcripts), defense (69) and detoxification (37 transcripts) are interrelated, in that pathogen, chemical (including internally generated and external toxins), or physiochemical insults induce a physiological stress, resulting in a defense or detoxification response. The stress category transcripts are primarily scored here as abiotic, including temperature, light and water. The stress transcripts are largely present in clusters 2 (16%), 4 (32%), 5 (26%) and 6 (32%), which have distinctly different temporal patterns of accumulation; these transcripts are not present in clusters 3, 8 and 9. Transcripts encoding proteins for defense can be categorized by response to biotic and abiotic insults; abiotic are generally grouped into the stress category. In this organization, defense responses against biotic insults are interpreted to be associated with annotations specifying fungal and bacterial pathogens, plus lectin binding. The biotic defense grouping represents 31 transcripts, with none present in clusters 3 and 8 and 64% in clusters 4, 5 and 7 wherein the initial transcriptional upregulation reflected in cluster 4 is opposite of the initial downregulation in clusters 5 and 7, implying different regulatory pathways and physiological roles. The temporal patterns in clusters 5 and 7 indicate the response are clearly (cluster 7) and possibly (cluster 5) influenced by *N. punctiforme*. Plant ubiquitous germin-like proteins (GLP) express both biotic and abiotic defense responses but here we emphasized the biotic potential. Twenty-one transcripts encoding GLP were differentially transcribed; GLP transcripts are not present in clusters 1, 8 and 9 and 81% are in clusters 5 and 7. The four interspecies interactions transcripts encode proteins with lectin binding domains (2 transcripts), isoflavone synthesis and leucine-rich repeats. Three of the transcripts are in cluster 2 and the other is in cluster 3; the transcriptional patterns in these clusters are similar in upregulation by the presence of *N. punctiforme* and differ by the timing of the upregulation. The detoxification category includes 31 transcripts and most prominent are peroxidases (16 transcripts) and glutathione S-transferase (5 transcripts, plus 1 glutamate-cysteine ligase). Transcripts encoding glutathione metabolism are present in clusters 1, 3, 4, 5 and 6, which showed mixed temporal patterns. Peroxidases are present in all clusters except 1, 8 and 9 and, also showed mixed patterns of accumulation, which indicates multiple transcriptional regulatory pathways in both detoxification subgroups. The broad temporal distribution of peroxidase encoding transcripts may ensure that peroxide does not accumulate under changing stress conditions during early N-starvation through to $N_2$-derived ammonium complementation.

Protein metabolism includes protein synthesis (24 transcripts), modification (44 transcripts) and turnover (51 transcripts). Protein synthesis transcripts are primarily ribosome biogenesis and translation processes; the transcripts are present in all clusters except 3 and 7, indicating both up- and downregulation of expression, with 67% of the transcripts in cluster 2, where the temporal patterns most obviously differ in the absence of *N. punctiforme*, and cluster 4, where the patterns are similar. Protein modification includes proteins catalyzing phosphatase, methyl transfer, glycosylation, isomerization, and protein-protein interactions. There are 16 chaperone transcripts that were present in all clusters except 1 and 8, with 44% of the transcripts in cluster 7, implying recovery of chaperones to the time 0 level occurs most robustly in the presence of *N. punctiforme*. Transcripts encoding peptidases and proteases involved in turnover were present in all clusters, indicating a mixed transcriptional response. Ubiquitin ligase and transfer activities represent 57% of the transcripts and were enriched in clusters 1, 5 and 7 where the presence of *N. punctiforme* had a negative (cluster 1), neutral (cluster 5) or positive (cluster 7) effect on transcriptional upregulation, perhaps implying targeting of specific proteins.

Approximately 9% of the differentially transcribed N-starvation genes encode transporters. Transcripts associated with a variety of types of transporters and their substrates are represented: 37 encode ion transporters, 16 encode ATP Binding Cassette (ABC) systems, only 3 encode major facilitator superfamily proteins, 13 are for efflux/export/secretion processes, 6 transcripts encode specific auxin transport (3) or efflux (3) proteins, 5 encode sugar transporters and 4 are aquaporin specific. While all clusters have at least two transcripts encoding a transport protein, the majority are broadly present in clusters 1 to 7, with each containing from 12, 14 or 18 transcripts. The 12 transcripts in each of clusters 1 and 4 reflect immediate upregulation; they differ by the repressive effect of the presence of *N. punctiforme* in cluster 1 and the decline in transcripts with time in both clusters in the presence and absence of *N. punctiforme*. Clusters 2 and 3 both have 14 transcripts, which display different patterns of initial downregulation in the absence of *N. punctiforme*, but upregulation in its presence, varying by the time of upregulation and by persistence of transcript accumulation. Clusters 5 and 6 also contain 14 transcripts each, with substantial differences in temporal patterns. The presence of *N. punctiforme* repressed transcription in cluster 6, while it paralleled the delayed upregulation seen in its absence in cluster 5. There are 18 different transcripts in cluster 7; in this case, transcription was severely depressed during N-starvation in the absence of *N. punctiforme*, but its presence increased transcription after about 10 days, implying recovery of steady state growth conditions. Specific transporters with relevance to N-limitation and excess, and symbiotic interaction are presented in the text.

Signal transduction is represented by 88 DEGs. These transcripts include 75 encoding protein kinases, of which 28 are serine/threonine (ST) kinases, 19 are for receptor kinases (RK), 18 are undefined active site of generic protein kinases (PK), 5 for tyrosine (TK) and 3 for histidine kinases (HK). Three guanine nucleotide-binding proteins (G-proteins) and 2 small G-proteins were also differentially expressed. Transcripts encoding signal transduction proteins sorted into all clusters but are more highly represented in clusters 2 (18 transcripts), 3 (10 transcripts), 4 (14 transcripts) and 7 (27 transcripts). Clusters 2, 3 and 7 reflect initial transcriptional downregulation which persisted in the absence of *N. punctiforme*, but its presence resulted in concurrent (cluster 2) or delayed (clusters 3 and 7) upregulation. Clusters 2 and 6 each contain transcripts for a G-protein and a small G-protein; the temporal patterns are almost reciprocal and imply the pair in cluster 6 are important to *A. punctatus* starved in the absence of *N. punctiforme* while those in cluster 2 are immediately complemented by its presence. The transcriptional regulatory mechanisms will likely differ in the two cases. In general, signal transduction proteins appear to be more involved in steady state symbiont-free and symbiotic growth than in an initial response to the stress of N-starvation.

Hormones, growth, and development is an interrelated category consisting of 68 transcripts. The N-starved *A. punctatus* differential transcriptome encodes 1 cytokinin in cluster 2; 3 gibberellic acid (GA) methyltransferases in clusters 2, 4 and 7; 4 GA signaling (alpha beta hydrolase domains) with 3 transcripts in clusters 4 and 1 in cluster 5; 5 auxins (indole-3-acetic acid-amido synthetase; IAA) in clusters 2, 4, 5 and 7 (2 copies); and 7 for jasmonic acid (JA), 6 of which encode 12-oxophytodienoate reductase in clusters 1, 3, 5 and 6, of which 2 copies are present in clusters 3 and 6. Overall, clusters 3 and 6 are enriched in IAA and JA synthesis; clusters 3 and 6 display reciprocal temporal patterns of expression. GA synthesis and signaling are in upregulated and slow decline cluster 4 which is unaffected by the presence of *N. punctiforme*. Growth and developmental processes operate either concurrently or as a continuum.

Growth related genes are scored as two groups: growth with 10 transcripts that are present in clusters 2 (2 transcripts), 3 (3 transcripts), 4 (1 transcript), 5 (2 transcripts) and 7 (2 transcripts); and growth plus general or vegetative development with 9 transcripts sorting in clusters 3 (1 transcript), 5, (1 transcript), 6 (5 transcripts) and 7 (2 transcripts). Specific developmental processes are annotated in the context of seed-producing vascular plants, of which there are 26 differentially expressed transcripts: 4 transcripts involved in stages of embryogenesis (clusters 1, 4, 6 and 7), 3 for gametophytes (all in cluster 5), 1 in shoot and root outcomes (cluster 1), leaves with 2 transcripts (clusters 3 and 4), flowers with 4 transcripts (clusters 1, 2 with 2 representatives, and 4), pollen with 5 transcripts (clusters 3, 7 and 9, with duplicates in cluster 4), stigma with 1 transcript (cluster 4), and seed formation and germination with 5 transcripts (clusters 2, 4 and 5 with 3 representatives). Two of the seed-related transcripts in cluster 5 encode proteins in the pirin family of transcriptional cofactors related to apoptosis.

The genome of *A. punctatus* potentially encodes approximately 620 transcription factors (TF), about 52% of which are not annotated in any described family (Li *et al.*, 2020). The N-starved differential transcriptome is represented by 36 genes encoding proteins for the transcription process, 29 of which are classified as TF. The non-TFs include 1 transcript encoding a protein of the preinitiation complex (cluster 3), 2 for RNA polymerase II activity (clusters 1 and 5), 1 of a sigma subunit (cluster 1), 1 for the elongation process (cluster 2), 1 for mRNA export (cluster 1) and one of a riboswitch (cluster 5). Of the 29 TF transcripts, 10 have no specific process association other than general TF and are found in clusters 1, 3, 4, 6, and 9; cluster 1 has 4 representatives and cluster 4 has 3, the remainder have 1 each). The remaining 19 TFs are associated specific processes or families such as the hormones auxin (1 in clusters 9), brassinosteroid (1 in cluster 7) and ethylene (2 in cluster 4); the nuclear factor kappa B (1 in cluster 7); DNA binding in the NAC family (1 in cluster 7); a pentatricopeptide repeat-containing activator (1 in cluster 4), an Fe supply factor (1 in cluster 2); a non-specific corepressor (1 in cluster 7); present are 4 zinc-finger-containing TFs, 1 in the WRKY53 family (cluster 5), another in the BBX14 family (cluster 3) and 2 are non-specific (clusters 2 and 4); the GRAS family is represented by 6 transcripts with 3 in cluster 5, 2 in cluster 1 and another one in cluster 6.

Oxygenic photosynthesis overwhelmingly defines the carbon and energy trophic level of plants but can be detrimental to plants under N-stress. The differential transcriptome has 35 genes encoding proteins indirectly or directly related to photosynthetic mechanism. Chloroplasts are directly related to photosynthesis as the cellular site of activity and represented by 17 nuclear-encoded transcripts which are distributed in all clusters except 8 and 9. Their functions include: carbon assimilation with the RuBisCO small subunit in cluster 3 and a carbonic anhydrase in cluster 7; carotenoid synthesis in clusters 1, 5 and 6; light induced chlorophyll synthesis in cluster 4; developmental aspects in cluster 4; single stranded DNA binding in cluster1; envelope membrane in cluster 3; oxidative stress in cluster 5; a protein of unknown function in cluster 2; a carbohydrate kinase involved in transcription in cluster 5 and 4 transport proteins with 1 in cluster 1 and 3 in cluster 3. There are 16 transcripts of genes encoding light harvesting complexes (LHC; 10 transcripts) and reaction center (6 transcripts) proteins all in cluster 3. Chloroplast and photosynthetic transcripts are most highly present in cluster 3 (22 transcripts), which were initially downregulated and then upregulated in the presence of *N. punctiforme* in a temporal pattern corresponding the production of $N_2$-derived ammonium.

Reference

**Li F-W, Nishiyama T, Waller M, *et al.*** 2020. *Anthoceros* genomes illuminate the origin of land plants and the unique biology of hornworts. Nature Plants **6**, 259–272.

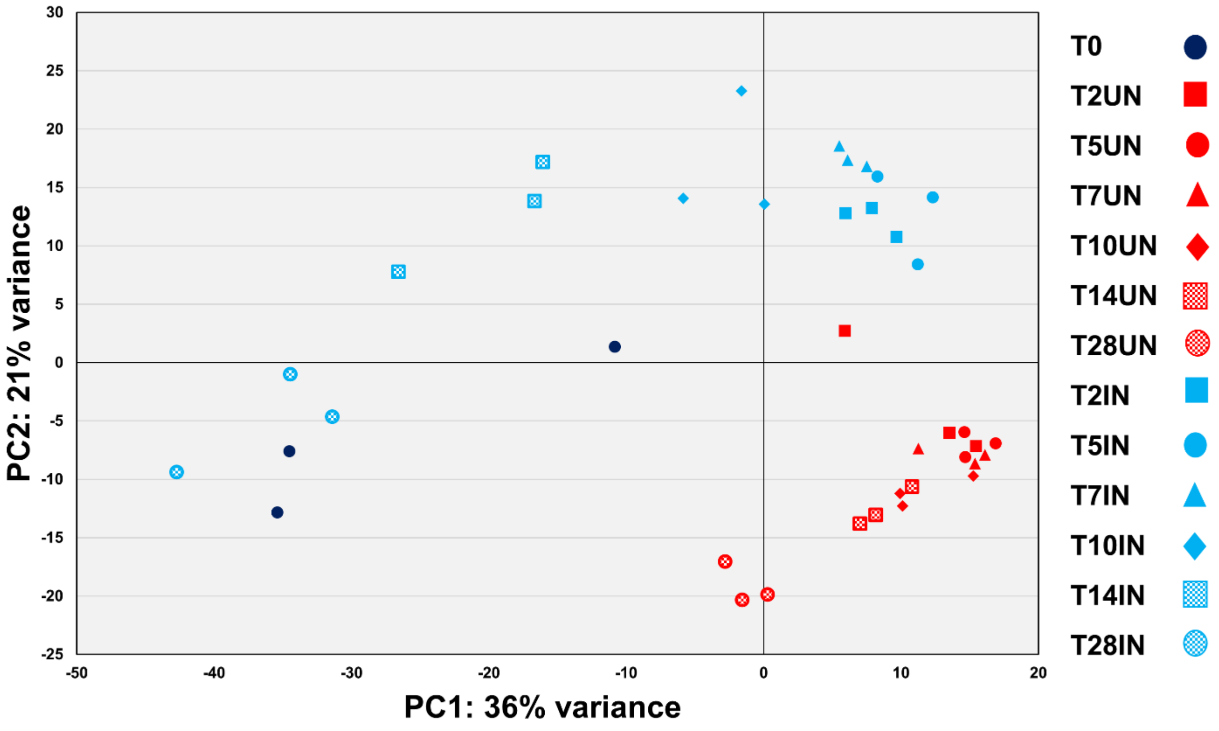


**Supplementary Figure 1**. PCA analysis of biological replicates.

Control (T0), cocultured (T2IN, T5IN, T7IN, T10IN, T14IN and T28IN) and symbiont-free (T2UN, T5UN, T7UN, T10UN, T14UN and T28UN) *A. punctatus* under N-starved conditions during time course RNA-Seq analysis.

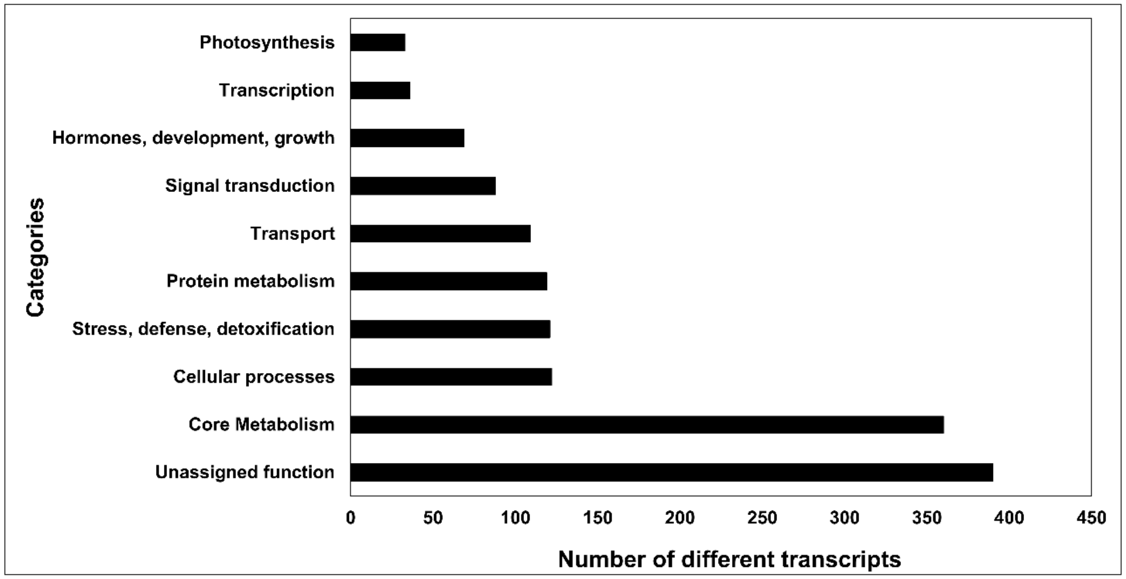


**Supplementary Figure 2.** Manually assigned metabolic categories of differentially transcribed genes observed in cluster analysis.
